# *Summix:* A method for detecting and adjusting for population structure in genetic summary data

**DOI:** 10.1101/2021.02.03.429446

**Authors:** IS Arriaga-MacKenzie, G Matesi, S Chen, A Ronco, KM Marker, JR Hall, R Scherenberg, M Khajeh-Sharafabadi, Y Wu, CR Gignoux, M Null, AE Hendricks

## Abstract

Publicly available genetic summary data have high utility in research and the clinic including prioritizing putative causal variants, polygenic scoring, and leveraging common controls. However, summarizing individual-level data can mask population structure resulting in confounding, reduced power, and incorrect prioritization of putative causal variants. This limits the utility of publicly available data, especially for understudied or admixed populations where additional research and resources are most needed. While several methods exist to estimate ancestry in individual-level data, methods to estimate ancestry proportions in summary data are lacking. Here, we present Summix, a method to efficiently deconvolute ancestry and provide ancestry-adjusted allele frequencies from summary data. Using continental reference ancestry, African (AFR), Non-Finnish European (EUR), East Asian (EAS), Indigenous American (IAM), South Asian (SAS), we obtain accurate and precise estimates (within 0.1%) for all simulation scenarios. We apply Summix to gnomAD v2.1 exome and genome groups and subgroups finding heterogeneous continental ancestry for several groups including African/African American (∼84% AFR, ∼14% EUR) and American/Latinx (∼4% AFR, ∼5% EAS, ∼43% EUR, ∼46% IAM). Compared to the unadjusted gnomAD AFs, Summix’s ancestry-adjusted AFs more closely match respective African and Latinx reference samples. Even on modern, dense panels of summary statistics, Summix yields results in seconds allowing for estimation of confidence intervals via block bootstrap. Given an accompanying R package, Summix increases the utility and equity of public genetic resources, empowering novel research opportunities.

## Introduction

Genetic summary data is a cornerstone of modern analyses. Allele frequencies (AFs) from publicly available data such as the genome Aggregation Database (gnomAD)^1,2^ and Allele Frequency Aggregator (ALFA) from dbSNP^3^ can be used to prioritize putative causal variants for rare diseases and as pseudo controls in case-control analysis ^4–7^. Compared to individual-level data, genetic summary data often has fewer barriers to access promoting open science and the broad use of valuable resources. However, summary-level genetic data frequently contain fine-scale and continental-level population structure. For instance, unquantified continental ancestry exists in gnomAD’s African/African American, American/Latinx, and Other groups as well as in other publicly available data (e.g. BRAVO server for TopMED)^2,8^. Using public data without accounting for the underlying population structure can lead to confounded associations and incorrect prioritization of putative rare causal variants.

The use of mixture models to estimate population structure has a history going back over two decades beginning with IMMANC ^9^ and later the commonly-used STRUCTURE^10^, originally using Dirichlet priors for multinomial modeling. Inference was performed via Markov Chain Monte Carlo, which limits tractability to datasets with thousands to tens of thousands of markers. As datasets grew with the advent of genome-wide arrays, and later with sequencing, new methods were designed with improved convergence characteristics such as the maximum-likelihood methods FRAPPE ^11^ and Admixture ^12^, as well as the variational method FastSTRUCTURE ^13^. Along the way, methods were developed to leverage pooled data^14^, e.g. iAdmix ^15^, with improvements to enhance supervised analysis in the Admixture framework ^16^. However, no method was designed explicitly and efficiently to model mixtures with genome-wide, summary statistic data that is common in modern genomics.

Individuals and samples from understudied or admixed populations are most likely to lack large public resources with precisely matched ancestry data. As a result, researchers and clinicians working with these populations are often left with a suboptimal choice: use the closest, but still poorly matched ancestral group or do not use the publicly available and highly useful resource ^7^. The former has the potential to produce biased results in the very populations where additional high-quality research is needed, while the latter is likely to suffer from smaller sample sizes and thus a loss of statistical power. This choice exacerbates inequities in research on understudied and admixed populations ^17,18^. These issues are magnified in the context of precision medicine where genetic summary data will likely not be sufficiently matched for the majority of people who themselves are a mixture of continental or fine-scale ancestries. Thus, methods to estimate and adjust for population structure within publicly available genetic summary data are needed.

Here, we present Summix, an efficient method that identifies, estimates, and adjusts for the proportion of continental reference ancestry in publicly available summary genetic data. We demonstrate the effectiveness of Summix in over 5,000 simulation scenarios and in gnomAD v2.1, including the ability to produce ancestry-adjusted AFs to tailor analyses to less-studied populations. Ultimately, Summix, and the accompanying R, Python, and Shiny app software, help to increase the efficacy and, importantly, the equity of valuable publicly available resources, especially for understudied and admixed samples.

## Material and Methods

### Summix

#### Estimating ancestry proportions

An observed single nucleotide polymorphism (SNP) AF can be described as a mixture of AFs across unobserved subgroups (e.g. continental ancestral populations). We estimate the group-specific mixing proportions, *π*_*k*_, by minimizing the least-squares difference between vectors of N SNPs for the observed AF, *AF*_*observed*_, and the AF generated from a mixture of K reference ancestry groups, *AF*_*ref, k*_, as shown in Equation (1).

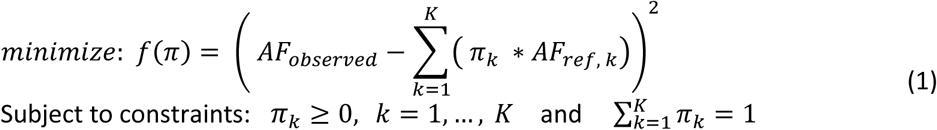

This objective function is quadratic and, as such, is continuous, convex, and easily differentiable; the inequality constraints are linear. Hence, a feasible minimizer will fulfill the Karush-Kuhn-Tucker (KKT) conditions for optimality. We use Sequential Quadratic Programming (SQP)^19,20^ a gradient-based, iterative algorithm for constrained, nonlinear optimization to efficiently estimate the proportion of each reference group. We obtain confidence intervals for the continental ancestry proportions using block bootstrapping as described below.

#### Ancestry-adjusted allele frequencies

Using estimated continental ancestry proportions, we update the AFs in the observed data matching the continental ancestry proportions of an individual or sample as follows in Equation (2). To estimate the ancestry-adjusted AF, K-1 homogenous reference ancestries are used. The ancestry not used is indexed as *l*. In theory, ancestry *l* can be any of the non-zero reference ancestry groups. Here, we choose *l* to be the most common ancestry present in the summary data.

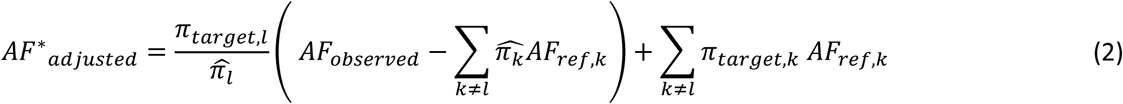

*Where*,

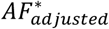 *is the ancestry-adjusted allele frequency*

*l - ancestry group for which the reference allele frequency data is not used*

*k - ancestry group index*

*π*_*target,k*_ *- population k ancestry proportion for target individual or sample*

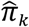-*estimated ancestry proportion of population k for observed publicly available summary data*

*AF*_*observed*_ *-allele frequency for observed publicly available summary data (e*.*g. gnomAD)*

*AF*_*ref,k*_ *-reference allele frequency for ancestry k*; K-1 homogenous reference ancestries are used.

Equation (2) can be used to estimate ancestry-adjusted AF for a homogenous population or an admixed population with given ancestry proportions. We evaluate both scenarios as described below.

### Data

All data were on genome build GRCh37.

#### 1000 Genomes and Indigenous American data

We used four continental ancestry groups from 1000 Genomes v5 phase 3 20150502 ^20^, African (AFR), East Asian (EAS), non-Finnish European (EUR), and South Asian (SAS), and an Indigenous American (IAM) sample for the reference data. The IAM Affymetrix 6.0 data were downloaded from ftp.1000genomes.ebi.ac.uk/vol1/ftp/technical/working/20130711_native_american_admix_train (data accessed October 2018) and had been previously harmonized with the 1000 Genomes data ^21^. 1000 Genomes AF were calculated as previously described ^22^. IAM AFs were calculated using PLINK 1.9. We excluded the admixed populations from the 1000 Genomes AFR continental ancestry: Americans of African Ancestry in SW USA (ASW) and African Caribbeans in Barbados (ACB). We merged the 1000 Genomes and IAM data keeping the subset of SNPs in both datasets. We limited further to biallelic non-palindromic SNPs with minor allele frequency (MAF)>1% in at least one continental ancestral group resulting in 613,298 SNPs.

#### gnomAD v2.1

Variants from gnomAD v2.1 (data accessed April 2019) were limited to biallelic and PASS resulting in 13,742,683 and 196,606,976 SNPs in the exome and genome gnomAD samples respectively. After limiting further to MAF >1% in at least one gnomAD group and merging with the reference data, 9,835 and 582,550 SNPs remained in the exome and genome data respectively.

#### ClinVar

ClinVar VCFs (GRCh37/hg19) were downloaded on September 25, 2020 from ftp.ncbi.nlm.nih.gov/pub/clinvar/vcf_GRCh37/. GnomAD v2.1 AF and ClinVar variants were merged by chromosome, base pair, and alleles. For three variants, there were multiple ClinVar allele IDs in the same position with the same alleles. All duplicate positions were retained. After merging and restricting to ClinVar classifications of “Uncertain Pathogenicity”, and “Conflicting reports of pathogenicity”, 42 and 122 variants were present in the exome and genome data, respectively. Ancestry-adjusted AFs were estimated for an African sample from the African/African-Amercian gnomAD control AFs using gnomAD EUR as the reference sample.

The American College of Medical Genetics (ACMG) guidelines^23^ designate >5% MAF in a control population as stand-alone strong evidence of a variant having benign impact for a rare Mendelian disorder. As such, for the classifications of “Uncertain Pathogenicity”, and “Conflicting reports of pathogenicity”, we identified variants where the ancestry-adjusted AF differed from the unadjusted AF with respect to the MAF > 5% threshold. Additionally, we identified variants with the classification “Pathogenic”, “Likely_pathogenic”, or “Pathogenic/Likely_pathogenic” with either adjusted or unadjusted AF above 5%.

### Simulations

Using the 1000 Genomes as reference data, we simulated SNP genotypes for all combinations and subsets of the five continental populations. SNP genotypes were simulated using the *rmultinom* R function with probability defined from the AFs for the continental reference ancestral populations assuming Hardy-Weinberg Equilibrium. Ancestry proportions were chosen randomly within an assigned proportion bin to ensure coverage across the range of possible values especially at the edges of the distribution: 0-0.015, 0.010-0.055, 0.05-0.105, 0.10-0.255, 0.25-0.505. The simulated proportion for the K^th^ ancestry group was chosen so that the ancestry proportions summed to one.

Simulating across all combinations of continental ancestry groups and the ancestry proportion bins resulted in 5,360 simulation scenarios. We used 1000 replicates within each simulation scenario to assess accuracy and precision. For each simulation replicate, we randomly sampled 100,000 SNPs. We define accuracy as the difference between the mean estimated ancestry proportion and simulated ancestry proportion. We define precision as the standard deviation of the simulation replicates. The simulation code is provided on the method’s GitHub site: https://github.com/hendriau/Summix_manuscript.

### Real Data Application

#### Estimating ancestry proportions

Continental reference ancestry proportions were estimated using the *summix* R function for gnomAD v2.1 African/African American, American/Latinx, East Asian, Other, Non-Finnish European, and South Asian exome and genome including all subgroups (e.g. controls, non-neuro). To estimate the ancestry group proportions, the freezes of 9,830 and 582,421 SNPs in the exome and genome samples respectively were used. To assess stability of the estimates over different numbers of SNPs, we estimated the ancestry proportions from 1000 random samples of sets of N SNPs: 10, 50, 100, 500, 1000, 2500, 5000, 10k, 50k, 100k for genomes, and 10, 50, 100, 500, 1000, 2500, 5000, 7500, 9000 for exomes. Finally, estimates of continental ancestry proportions from Summix using summary AF data were compared to estimates from ADMIXTURE ^12^ using individual-level data for a sample of N=85 unrelated individuals from the Peruvian 1000 Genomes data.

#### Block bootstrapping

We used block bootstrapping to estimate uncertainty for the ancestry proportion estimates. We used the sex-averaged centimorgan (cM) map created from Bherer et al ^24^ to define 1 cM blocks throughout the genome. The *na*.*approx* function ^9,25^ from the ***zoo*** R package ^25^ was used to linearly interpolate cM for SNPs in our dataset that were not observed in Bherer et al. This resulted in 3357 1cM blocks across the genome. Five and 129 SNPs in the exome and genome data respectively that were outside the genetic regions contained in the Bherer et al data set were not linearly interpolated. This resulted in 9,830 and 582,421 SNPs in 2206 and 3353 1cM blocks for the exome and genome gnomAD samples respectively. This final sample was used for all real data analysis. We used 1,000 bootstrap replicates to estimate 95% block bootstrap confidence intervals. The lower and upper confidence intervals were estimated from the 2.5 and 97.5 percentiles of the block bootstrap distribution.

#### Estimating ancestry-adjusted allele frequencies

We estimated and assessed ancestry-adjusted AFs for two samples: (1) an African sample (100% African) estimated from the African/African American gnomAD AFs and (2) an admixed Peruvian sample with average ancestry proportions of 74.6% Indigenous American, 19.9% European, 2.8% African ancestry, 2.7% East Asian ancestry estimated from the American/Latinx gnomAD AFs. The continental ancestry proportions for the admixed Peruvian population were estimated from a subset of unrelated individuals (N = 85) from the 1000 Genomes Peruvian sample using ADMIXTURE ^12^. For both the African sample and the Peruvian sample, ancestry-adjusted AFs were estimated using reference AF from either 1000 Genomes or gnomAD. We used reference groups with ≥2% estimated ancestry proportion in the observed gnomAD v2.1 group and normalized the estimated ancestry proportions to total 1. This resulted in K=2 ancestry groups for the African/African American group and K=4 for the American/Latinx group. For the African/African American group with K=2 reference groups, non-Finnish European reference AFs were used (excluding African reference). For the Peruvian population where K=4, non-Finnish European, African, and East Asian reference AFs were used (excluding Indigenous American reference). To estimate the Peruvian ancestry-adjusted AFs using gnomAD reference populations, ancestry-adjusted AFs for a 100% African sample were estimated from the gnomAD African/African American reference group.

To assess the accuracy of the ancestry-adjusted estimates, we compared the ancestry-adjusted and unadjusted gnomAD AFs to 1000 Genomes AFs for the target ancestral population (i.e. African or Peruvian Latinx). For these comparisons, we filtered out variants that were called in fewer than 25% of the gnomAD group. This removed 120 and 128 variants for African/African American and American/Latinx gnomAD exome groups and 8 and 11 variants for African/African American and American/Latinx gnomAD genome groups, respectively. We calculated both the absolute and relative difference as shown in Equation (3).

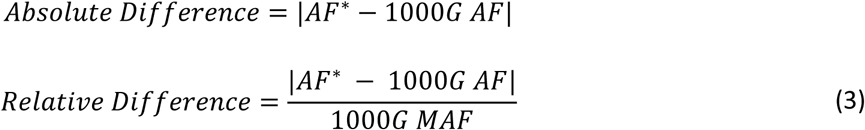

To test whether differences varied by adjustment group, we use a linear mixed effects model with SNP and cM block as random effects with the ***lmer*** function from the **lme4** package^26^. We tested all SNPs as well as within AF bins defined by the target 1000 Genomes reference ancestry (AF bin: <0.01, 0.01-0.02, 0.02-0.05, 0.05-0.1, 0.1-0.3, 0.3-0.5, 0.5-0.7, 0.7-0.9, 0.9-0.95, 0.95-0.98, 0.98-0.99, ≥0.99). Pairwise comparisons were adjusted for post-hoc multiple testing using Tukey adjustment. We assessed agreement between 1000 Genomes and ancestry-adjusted AFs or unadjusted AFs using Lin’s Concordance Correlation Coefficient (Lin’s CCC) ^27,28^ and 95% confidence intervals estimated using the *CCC* function in the ***DescTools*** R package version 0.99.38 ^29^.

### Reanalysis of *PADI3*

To demonstrate the utility of Summix’s ancestry-adjusted estimates in case-control analysis using external controls, we reanalyzed data from Malki et al ^7^. Malki et al. used gnomAD v2.1 African/African American as controls in a case-control analysis to show that *PADI3* is associated with Central Centrifugal Cicatricial Alopecia in women of African ancestry ^7^. We estimated the ancestry-adjusted AF for *PADI3* variants identified by Malki et al. for a sample with 100% African ancestry. Ancestry-adjusted allele counts (ACs) were estimated by multiplying the ancestry-adjusted AF by the variant specific allele number. The number of individuals in gnomAD v2.1 with at least one minor allele were calculated as the number of minor alleles minus the number of homozygotes. Using the case numbers reported from their manuscript, we repeated the case-control analysis using the original unadjusted gnomAD v2.1 data and the ancestry-adjusted gnomAD v2.1 results for a homogeneous African sample. As in Malki et al., we provide p-values for both a chi-square (*χ*^*2*^) test of independence and Fisher’s exact test.

### Software and code

Summix is available in R and Python. The Python (v3.7) package, ***summixpy***, contains two main Python scripts. The first function, *summix*.*py*, estimates the proportions of reference ancestry groups for an observed sample. The second function, *adjAF*.*py*, estimates the ancestry-adjusted AF. The package contains example genetic data and analysis presented in a Jupyter Notebook all of which are hosted, along with the package here: https://github.com/jordanrhall/summix_py.

The **Summix** R package enables both estimation of reference ancestry groups using the *summix* function and ancestry-adjusted AFs using the *adjAF* function. More details including example data and implementation are available in our package, which is hosted on Bioconductor and our GitHub site: https://github.com/hendriau/Summix.

### Shiny app

The Shiny app can be found here: http://shiny.clas.ucdenver.edu/Summix. Within the Shiny app, users can estimate and visualize ancestry proportions for three gnomAD ancestry groups (i.e. African/African American, American/Latinx, Other) for both the exome and genome data.

## Results

### Simulations

Summix achieved accuracy within 0.001% and precision within 0.1% across all simulation scenarios (**Tables S1-S5, Figures 1, S1-S4**). Accuracy of the proportion estimates was consistent across the range of simulated mixture proportions with a slight increase in bias near 0 and 1. While bias and variability in the estimates was small for all ancestral groups, AFR had the lowest variability followed by IAM, EAS, EUR, and then SAS with the highest variability across simulation replicates (**Tables S1-S5, Figures 1, S1-S4**). For simulation scenarios with non-zero proportions simulated for all five ancestry groups, the standard deviation was: SD_AFR_=4.31E-05, SD_IAM_=5.53E-05, SD_EAS_=7.58E-05, SD_EUR_=9.18E-05, SD_SAS_=1.21E-04. This trend was consistent across all simulation scenarios.

**Figure 1.**
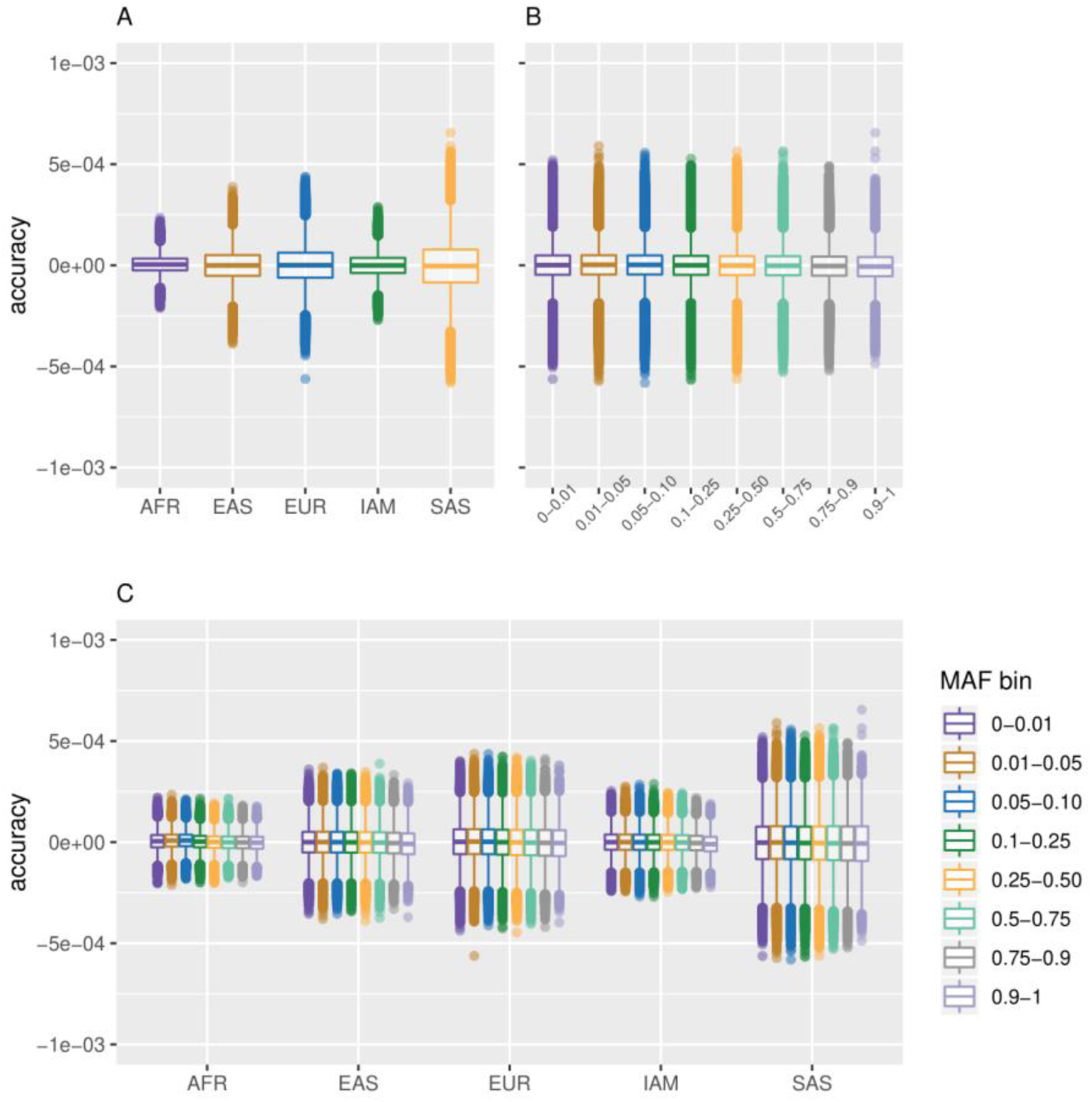
Simulation results for five ancestry parameters. Accuracy is defined as the absolute difference between the estimated ancestry proportions and given ancestry proportions within simulations. Five reference ancestries were used to simulate genotypes of an admixed population. **A)** Accuracy separated by ancestry. **B)** Accuracy separated by minor allele frequency. **C)** Accuracy separated by both ancestry and minor allele frequency.

### Application to gnomAD

#### Estimating ancestry proportions

We estimated the proportion of reference continental ancestry groups in gnomAD v2.1 African/African American, American/Latinx, East Asian, Other, Non-Finnish European, and South Asian groups and subgroups (e.g. controls, non-TopMED) for both the exome and genome data (**Tables 1, S6, S7**). As expected, the African/African American groups have primarily AFR (>80%) and EUR ancestry (∼15%) likely due to contribution from African American individuals. The exome and genome American/Latinx gnomAD groups had high proportions of both EUR and IAM ancestry (i.e. >35%) and ancestry proportion estimates between 1-6% for the other reference groups. Interestingly, >1% SAS ancestry was estimated in both the exome and genome American/Latinx gnomAD groups perhaps due to misspecification from a limited number of reference groups. The exome and genome Other gnomAD groups were primarily EUR reference ancestry (>77%). The estimated reference proportions for Non-Finnish Europeans and East Asian were very homogeneous with >96% EUR and 100% EAS estimated reference ancestries, respectively. The South Asian gnomAD exome group had 85% estimated SAS ancestry and ∼15% estimated EUR reference ancestry as expected due to the known ancient admixture events in the region^30–32^.

**Table 1.**
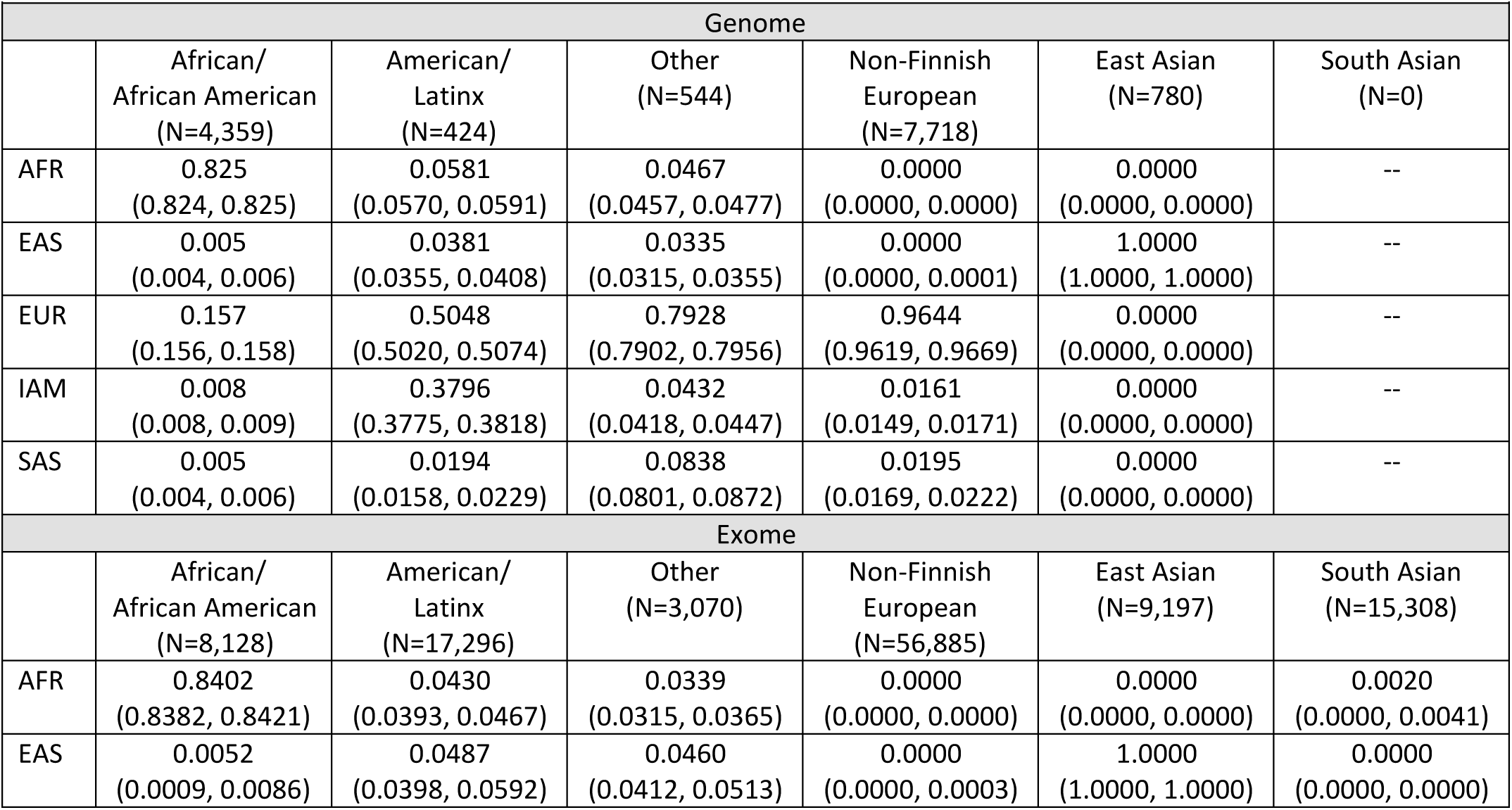
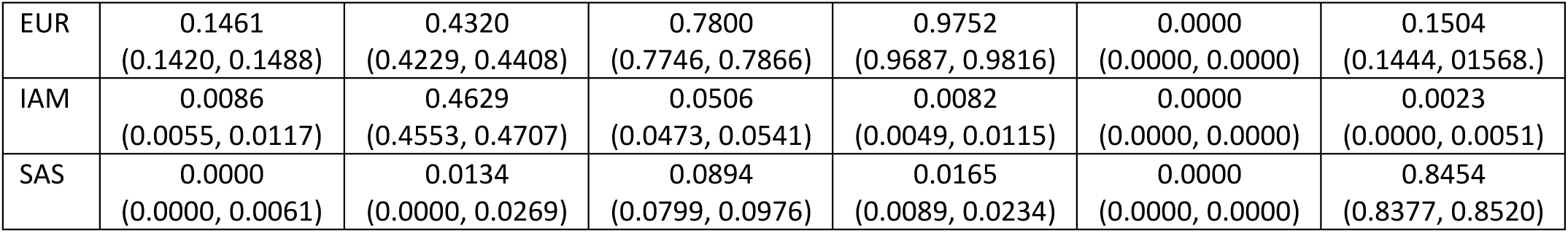
Estimated ancestry proportions for gnomAD groups (95% Block Bootstrap CI)

Despite large differences in sample sizes, the estimated proportion of reference ancestry groups was similar (i.e. within 2%) between exome and genome samples for all groups except American/Latinx where the exome and genome estimates differed by >5% for the EUR and IAM reference proportions. The reference ancestry proportion estimates for the gnomAD v2.1 subgroups (i.e. controls, non_cancer, non_neuro, and non_topmed) were mostly similar, within ∼2% of each other and of the overall gnomAD v2.1 group estimates. The exception was for the American/Latinx and Other genome groups, which sometimes varied by 5-10%. These groups had sample sizes (N<600) and thus were likely more susceptible to the inclusion or exclusion of subsamples of individuals. Complete results are shown in **Table S6-S7**.

We evaluated Summix’s ability to estimate reference ancestry proportions using smaller numbers of randomly chosen SNPs. We find that the ancestry proportion estimates stay unbiased regardless of the number of SNPs used to estimate ancestry while the precision decreases as the number of SNPs decreases. The precision is still fairly tight for down to ∼500 SNPs, especially when estimating the African/African American gnomAD samples (exomes: SD_AFR_=0.0039, SD_IAM_=0.0052, SD_EAS_=0.0054, SD_EUR_=0.0065, SD_SAS_=0.0067). This suggests that far fewer SNPs are likely needed to arrive at sample estimates of ancestry proportions (**Figure 2, Figure S5**).

**Figure 2.**
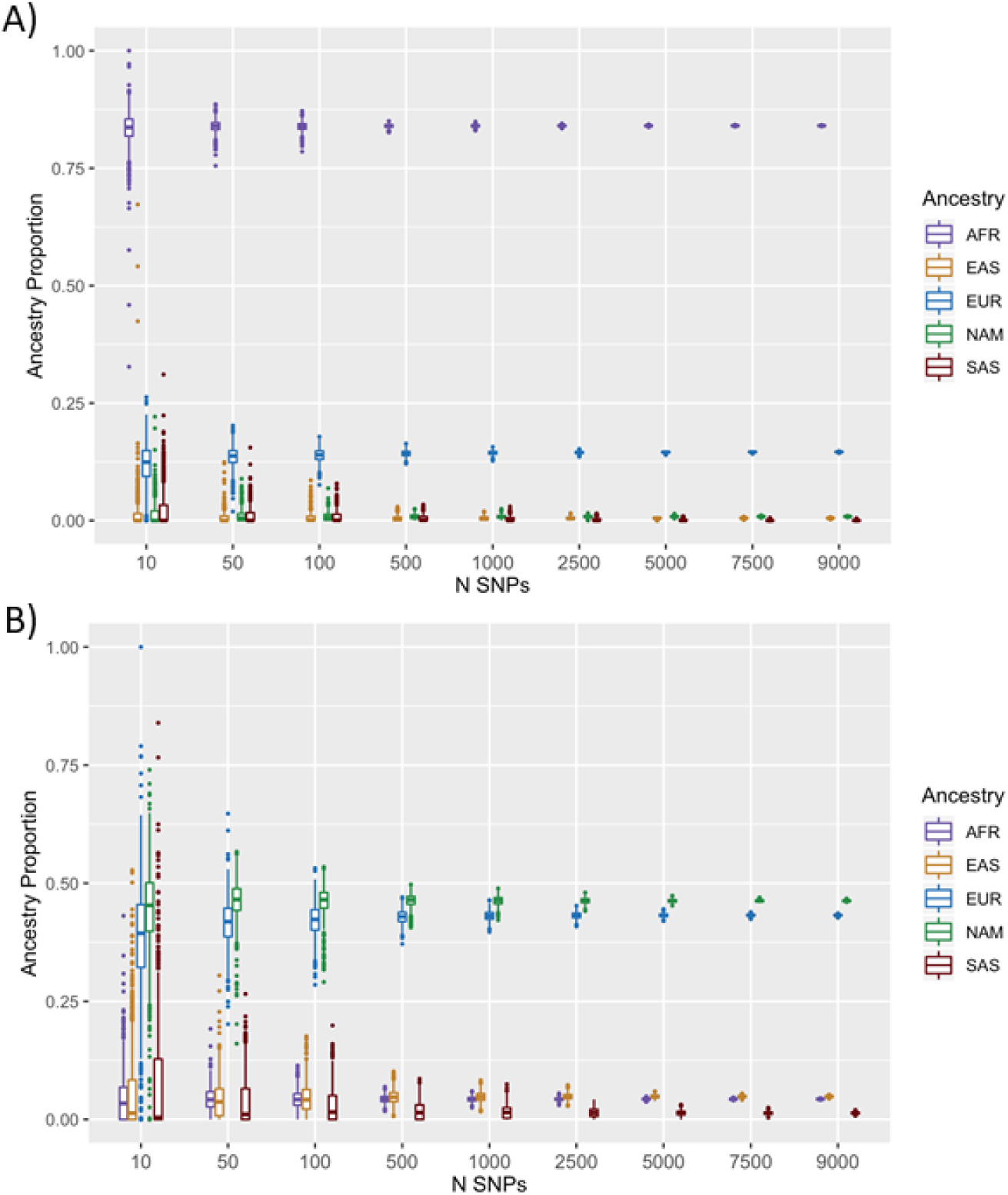
Precision in ancestry estimates for AFR and AMR gnomad groups by number of SNPs. Number of SNPs (x-axis), estimated ancestry proportion (y-axis) for 1,000 replicates; **A)** AFR exome. **B)** AMR exome.

#### Ancestry-adjusted allele frequencies

For both exome and genome data, we estimated the ancestry-adjusted AF for gnomAD African/African American for a target sample with 100% continental African ancestry and for gnomAD American/Latinx for a target Peruvian sample with 74.6% Indigenous American, 19.9% European, 2.8% African ancestry, 2.7% East Asian ancestry proportions. Compared to the unadjusted AFs, the ancestry-adjusted AFs had significantly smaller absolute and relative differences with the target group AF (**Table 2, Tables S8-S9**, p < 1E-16) regardless of reference data used (i.e. 1000 Genomes or gnomAD). The relative difference was greatest at small MAF while the absolute difference increased as AF increased, consistent with expectation. For SNPs with an alternative AF > 0.9 in 1000 Genomes, the absolute and relative difference between the unadjusted gnomAD and target 1000 Genomes AFs was especially large. This is likely due to both a relatively small number of SNPs in these groups and differences between 1000 Genomes samples and the reference genome. The absolute and relative differences in SNPs with AF > 0.9 is considerably reduced for the ancestry-adjusted AFs (**Figures 3-4, Tables 2, S8-S9**).

**Table 2.**
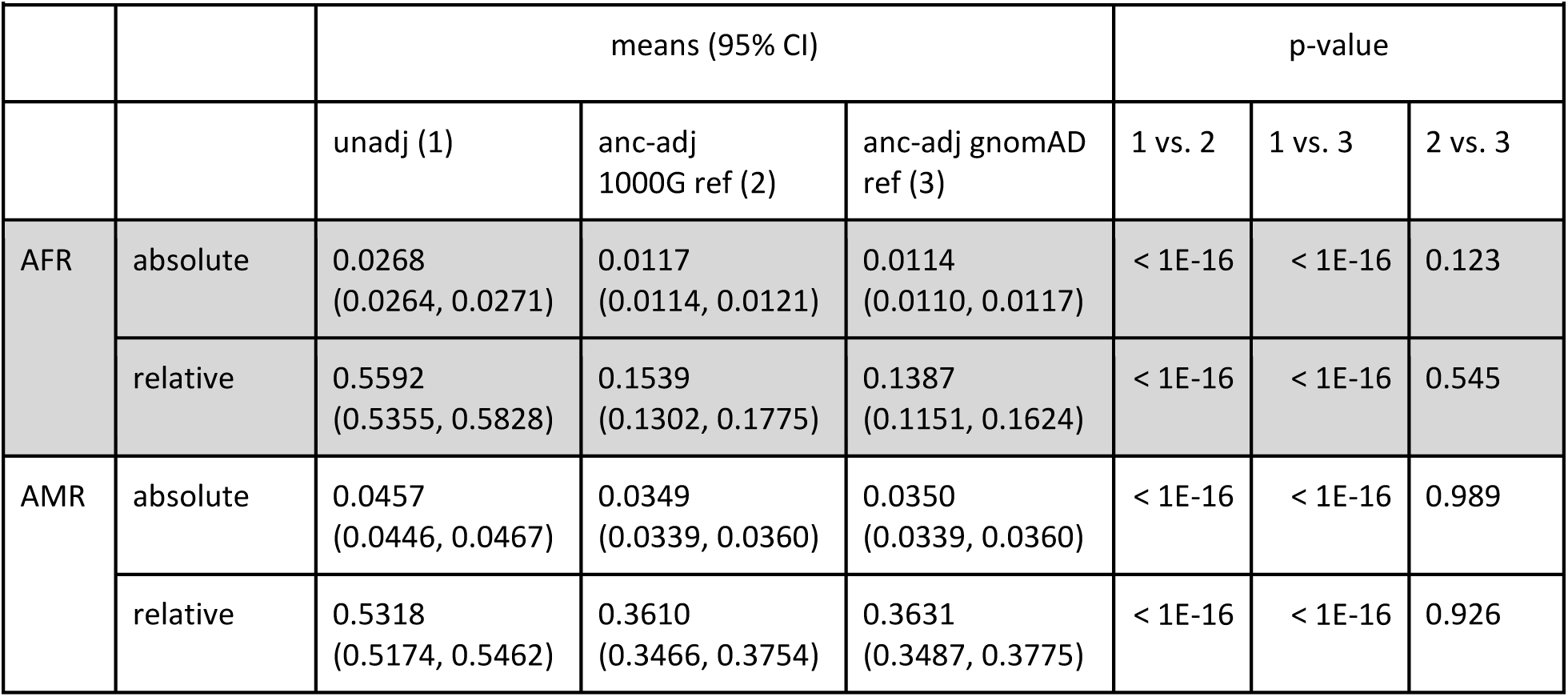
Absolute and relative difference between unadjusted and adjusted gnomAD AF and target sample in the exome data.

|  |  |  |  |  |  |  |  |
| --- | --- | --- | --- | --- | --- | --- | --- |
|  |  | means (95% CI) |  |  | p-value |  |  |
|  |  | unadj (1) | anc-adj<br>1000G ref (2) | anc-adj gnomAD<br>ref (3) | 1 vs. 2 | 1 vs. 3 | 2 vs. 3 |
| AFR | absolute | 0.0268<br>(0.0264, 0.0271) | 0.0117<br>(0.0114, 0.0121) | 0.0114<br>(0.0110, 0.0117) | < 1E-16 | < 1E-16 | 0.123 |
|  | relative | 0.5592<br>(0.5355, 0.5828) | 0.1539<br>(0.1302, 0.1775) | 0.1387<br>(0.1151, 0.1624) | < 1E-16 | < 1E-16 | 0.545 |
| AMR | absolute | 0.0457<br>(0.0446, 0.0467) | 0.0349<br>(0.0339, 0.0360) | 0.0350<br>(0.0339, 0.0360) | < 1E-16 | < 1E-16 | 0.989 |
|  | relative | 0.5318<br>(0.5174, 0.5462) | 0.3610<br>(0.3466, 0.3754) | 0.3631<br>(0.3487, 0.3775) | < 1E-16 | < 1E-16 | 0.926 |

**Figure 3.**
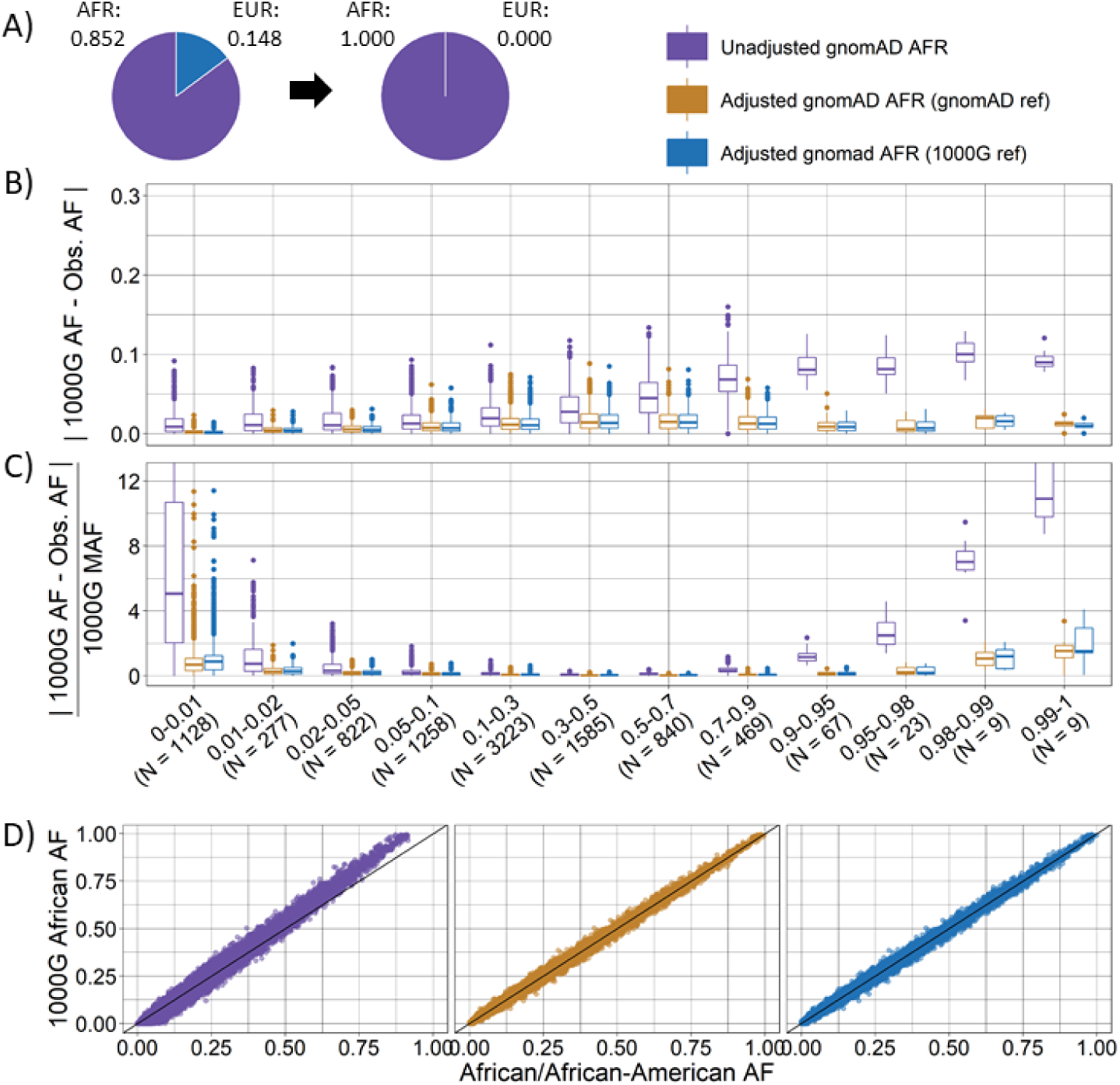
Ancestry-adjusted vs. unadjusted allele frequency for gnomAD African/African American exomes for a target sample with African ancestry. Ancestry-adjusted AF was estimated for a target sample with 100% African ancestry using gnomAD (orange) or 1000 Genomes (blue) Non-Finnish European as reference and compared to unadjusted AF (purple) for 9,710 SNPs. **A)** ancestry proportions for gnomAD African/African American exomes (AFR = 0.852, EUR=0.148) and target sample (AFR = 1); **B)** absolute difference between target sample AF (1000 Genomes African ancestry) and unadjusted or ancestry-adjusted gnomAD AF by 1000 Genomes AF category; **C)** relative difference between target 1000 Genomes African ancestry AF and unadjusted or ancestry-adjusted gnomAD AF by 1000 Genomes AF category; unzoomed figures B and C are available in the supplemental (**Figure S10**). **D)** scatter plot of target sample 1000 Genomes AF (y-axis) and unadjusted (left), ancestry-adjusted with gnomAD reference (center), and ancestry-adjusted with 1000 Genomes reference (right) gnomAD AF (x-axis).

**Figure 4.**
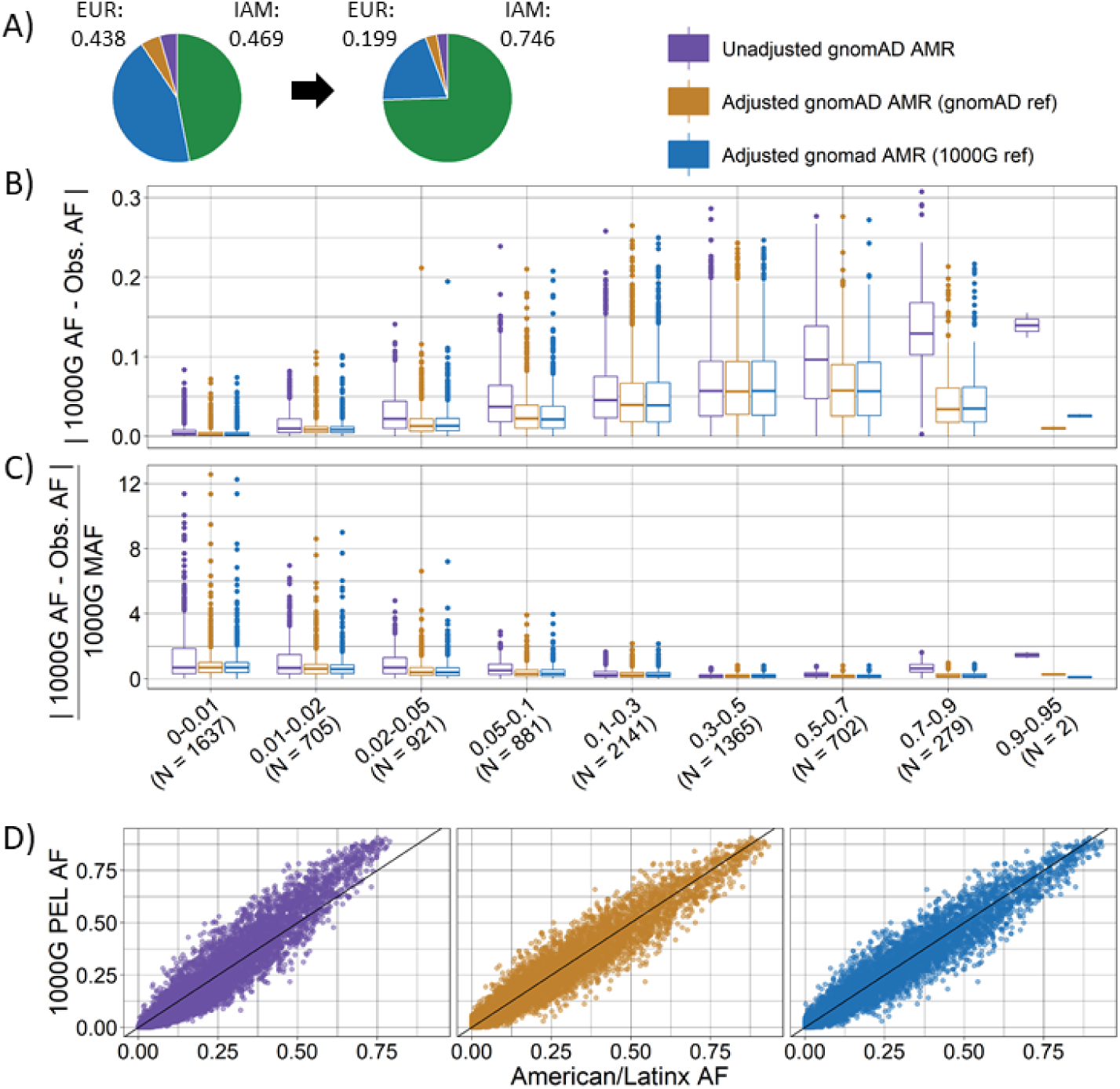
Ancestry-adjusted vs. unadjusted allele frequency for gnomAD American/Latinx exomes for a target sample of Peruvian ancestry. Ancestry-adjusted AF was estimated for a target Peruvian sample using gnomAD (orange) or 1000 Genomes (blue) East Asian, European, and African as reference ancestral populations and compared to unadjusted AF (purple) for 8,633 SNPs. **A)** normalized ancestry proportions estimated for gnomAD American/Latinx exomes (purple AFR = 0.044, orange EAS=0.049, blue EUR=0.438, green IAM=0.469) and target Peruvian ancestry proportions (purple AFR = 0.028, orange EAS=0.027, blue EUR=0.199, green IAM=0.746); **B)** absolute difference between target 1000 Genomes Peruvian ancestry AF and unadjusted or ancestry-adjusted gnomAD AF by 1000 Genomes AF category; **C)** relative difference between target 1000 Genomes Peruvian ancestry AF and unadjusted or ancestry-adjusted gnomAD AF by 1000 Genomes AF category; unzoomed figures B and C are available in the supplemental (**Figure S11**). **D)** scatter plot of target 1000 Genomes AF (y-axis) and unadjusted (left), ancestry-adjusted with gnomAD reference (center), and ancestry-adjusted with 1000 Genomes reference (right) gnomAD AF (x-axis).

The ancestry-adjusted AFs using 1000 Genomes or gnomAD as reference data were very similar. In the exome data, we observed no systematic significant absolute or relative differences between reference datasets (i.e. gnomAD reference vs. 1000 Genomes reference) with the exception of relative difference for the African 0-0.01 bin. Likely due to larger numbers of SNPs in the genome data, we do see significant, albeit very small, differences by reference ancestry between the ancestry-adjusted AF and the AFs in the target 1000 Genomes sample (**Tables S8-S9**). These differences, while statistically significant, were 10 to 100 times smaller compared to the unadjusted AF.

Lin’s CCC was used to estimate agreement between the target sample and the gnomAD AF. In gnomAD exomes, Lin’s CCC estimates were higher for the ancestry-adjusted compared to the unadjusted AFs regardless of reference data for both the African group (estimate [95% CI]: adj gnomAD_ref = 0.9977 [0.9976, 0.9978]; adj 1000G_ref = 0.9976 [0.9975, 0.9977], unadj = 0.9866 [0.9862, 0.9870]) and American group (adj gnomAD_ref = 0.9689 [0.9676, 0.9702]; adj 1000G_ref = 0.9691 [0.9678, 0.9704]; unadj = 0.9438 [0.9417, 0.9458]). We found similar results for the gnomAD genome data. Both the ancestry-adjusted and unadjusted AFs differ more for the gnomAD American/Latinx group compared to the target 1000 Genomes Peruvian AFs than for the African comparison. This is perhaps due to a larger number of reference ancestry groups or more heterogeneity in Latinx compared to African American samples.

Within the gnomAD genome data, we identified some SNPs with a large mismatch between 1000 Genomes and gnomAD AF (**Figures S6 – S9**). These SNPs appear to be mostly multi-allelic with one or more indels as the additional alleles **(Tables S10-S11**). We recommend caution when using these multi-allelic variants as more research and potentially external validation are likely needed to estimate the true AF of all alleles present.

### Summix vs ADMIXTURE Ancestry Estimates

We estimated ancestry proportions for the 1000 Genomes Peruvian sample of 85 unrelated individuals using AFs with Summix and individual-level data with ADMIXTURE. Assuming four reference ancestry groups, we estimate 0.033 AFR (0.032, 0.035), 0.035 EAS (0.031, 0.039), 0.209 EUR (0.206, 0.212), 0.723 IAM (0.719, 0.726) for Summix compared to 0.028 AFR, 0.027 EAS, 0.199 EUR, 0.746 IAM for ADMIXTURE.

### Reanalysis of *PADI3*

Summix can be used to estimate and adjust for ancestry in analyses that use gnomAD and other summary data as external controls. As an exemplar, we repeated the case-control analysis of *PADI3* from Malki et al ^7^. We found the p-values were higher for the ancestry-adjusted allele counts (ACs) (chi-squared p-value = 0.114, Fisher’s exact test p-value = 0.101) compared to the unadjusted gnomAD v2.1 African/African American ACs (chi-squared p-value = 0.029, Fisher’s exact test p-value = 0.031) (**Tables S12-S13**). As expected, this shows that association results are not robust to differences in ancestry. It is likely that the cases used by Malki et al ^7^ were not 100% African ancestry. Summix could be used to estimate ancestry-adjusted AC in gnomAD given the exact ancestry proportions in the cases. We expect the p-values of association would likely lie between the unadjusted and adjusted p-values provided given the proportion of African ancestry is likely between gnomAD unadjusted (0.852) and adjusted (1).

### ClinVar

As an exemplar for the potential utility of ancestry-adjusted AF in clinical settings, we compare the ancestry-adjusted AF for 100% African ancestry to the unadjusted AF for the gnomAD African/African American sample for a subset of Clinvar variants labeled as pathogenic, uncertain pathogenicity, or conflicting reports (**Methods**).

Based on previous ACMG guidelines^23^, we focused on variants labeled as either “Uncertain Pathogenicity”, and “Conflicting reports of pathogenicity.” We identified 68 unique ClinVar variants at 67 positions on the exome and genome (including 11 variants that were identified on both the exome and genome) where the ancestry-adjusted AF was above 5% and unadjusted AF was below 5%; and 42 variants at 41 positions where the ancestry-adjusted AF was below 5% and the unadjusted AF was above (5 of these variants were identified on both the exome and genome). Overall, we find minor differences of less than 0.05 between ancestry-adjusted and unadjusted AF. Eleven variants had differences greater than 0.05 in AF (**Table S14**). Some of these variants likely warrant further follow-up.

We identified 29 variants with unique ClinVar IDs (18 in the genome data, 2 in the exome data, and 9 in both the exome and genome data) listed as pathogenic or likely-pathogenic in ClinVAR with AF>5% for either the unadjusted or adjusted gnomAD African/African American samples respectively. All but three of these variants had adjusted and unadjusted AF>5% with most variants being very common (e.g. MAF>20%) (**Table S15**). Further inspection of these variants indicated little to no support for pathogenicity. Most of these (N=24) do not have assertion criteria provided and 13 were submitted to the ClinVar repository well before the 2015 update to the ACMG guidelines prompted by increased use of high-throughput sequencing^33^. These variants likely require further review. One of the variants identified as pathogenic and having a high AF is rs429358, one of two variants that defines the APOE-ε4 allele that infers an increased risk of Alzheimers. The high AF for rs429358 is expected. The APOE-ε4 allele confers increased risk of Alzheimer’s Disease in various ancestries including European^34^ and African^35,36^ although heterogeneity is observed in effect size and allele frequency across ancestries^37^.

## Discussion

Here we describe Summix, a fast, accurate, and precise method to estimate and adjust for population structure within publicly available genetic summary data. We evaluate Summix in over 5,000 simulation scenarios showing the accuracy and precision is within 0.001% and 0.1% respectively. In gnomAD, we find heterogeneous ancestry similar to what is expected in African/African American, American/Latinx, Other, and South Asian groups. We provide ancestry proportion estimates for all gnomAD groups and subgroups as well as ancestry adjusted AFs for an African sample and for a Peruvian sample for others to use.

Using the estimated proportion of continental ancestry groups, we produce ancestry-adjusted AFs for target samples with either continental African ancestry or Peruvian ancestry. When comparing to a sample with matching ancestry, we find that the unadjusted AFs differ significantly more than the ancestry-adjusted AFs regardless of reference ancestry data used (i.e. gnomAD vs 1000 Genomes). The African ancestry-adjusted AFs are more similar to the target 1000 Genomes African AFs than the Peruvian ancestry-adjusted AFs are to the target 1000 Genomes Peruvian sample. The increased dissimilarity for the Peruvian sample may be due to more than two predominant reference ancestry groups, or ancestral differences (including admixture) between gnomAD American/Latinx and 1000 Genomes Peruvian.

While the AFs of putative causal variants from a breadth of ancestral populations in public databases is useful for assessing evidence for clinical pathogenicity of a genetic variant, checking the AF in an ancestral sample that matches the ancestry of the person with the disease is most useful. Summix can be used to provide ancestry-adjusted AFs to precisely match ancestry increasing clinical utility of public datasets that may not have an ancestry that matches the patient. Additionally, Summix can produce ancestry-adjusted AFs matching the ancestry proportions for an external control sample. Here, we evaluate the potential utility of Summix by producing ancestry-adjusted AFs for ClinVar variants and for a case-control analysis of *PADI3*, a gene where gnomAD was used as external controls to identify association with Central Centrifugal Cicatricial Alopecia in women with African ancestry. While we find mostly minor discrepancies in the unadjusted and ancestry-adjusted AFs, we note that these differences can result in changed levels of evidence of association for case control analysis and for prioritization of putative causal variants for follow up. This emphasizes the importance of matching by or adjusting for ancestry differences whenever possible.

Here, we estimate genome-wide continental ancestry proportions. While the mean local ancestry proportions for a sample often approach genome-wide ancestry proportions ^38^, there may exist regions of the genome, e.g. regions of selection, where the local ancestry for the sample differs substantially from genome-wide ancestry proportions ^39,40^. We expect the ancestry-adjusted AF estimates may be less accurate in regions where the average local ancestry proportions differ from the genome-wide estimates. Our results suggest that Summix can estimate ancestry proportions accurately, although much less precisely, using a relatively small number of randomly chosen SNPs (e.g. ∼100). This suggests that Summix may be able to estimate the proportion of local continental ancestry. Here, we evaluate subsets of randomly chosen SNPs, albeit reflecting genome-wide coverage. It could be that using ancestry informative markers (AIMs)^41^ or removing uninformative markers could increase the precision of Summix further enabling the estimation of local ancestry proportions.

There are several drawbacks to our current method and implementation. First, our method is currently only able to estimate the proportion of provided reference ancestry groups. We recommend the user include all expected ancestral populations in the reference dataset. Second, here we only evaluate the ability of Summix to estimate five broad continental ancestries. We are actively working on an extension to identify and estimate the proportion of ancestry not in the reference data and are evaluating the performance of Summix on a broader reference panel including fine-scale ancestry. Lastly, differences in ancestry is not the only aspect of public databases that can cause confounding in analyses using external public controls. Differences in sequencing technology and computational variant calling pipelines can also cause biases in allele frequencies due to non-exchangeability of individuals. Many methods have been and are being developed to adjust for this bias when using external controls ^4–6,42,43^.

There are many extensions and applications for Summix beyond those evaluated here. First, Summix can be used with any reference ancestry data needing only AFs. While we provide ancestry-adjusted AFs for a sample, the ancestry-adjusted AFs could be used for individuals providing potential utility in the clinic. Summix has the potential to be applied to other summary datasets where AFs are provided or can be derived such as GWAS summary statistics, which are widely available online ^44^. Lastly, Summix has similarities to deconvolution methods used in single-cell and other ‘omics data types^45–47^, suggesting paths of future development and application.

We provide an R package, Python function, Shiny app, and GitHub site to encourage reproducibility, broad use, and further development of our method. We hope that the methods presented here will be used and extended to improve the utility of valuable publicly available resources especially for individuals and studies with admixed or understudied ancestry.

## Supplemental Data

Supplemental Data include eleven figures and fifteen tables.

## Acknowledgments

We thank the Education Through Undergraduate Research and Creative Activities (EURēCA!) program and Undergraduate Research Opportunities Program (UROP) through the University of Colorado Denver for supporting many of the undergraduate researchers on this project. This work was supported by the National Human Genome Research Institute (R35HG011293 and U01HG009080 to AEH and CGR; U01HG009080-05S1 to CGR).

## Data and Code Availability

Simulation data and code are available at https://github.com/hendriau/Summix_manuscript. Public data: gnomAD v2 data was downloaded from https://gnomad.broadinstitute.org/downloads on Oct. 11, 2018, 1000 Genomes data was downloaded from ftp://ftp.1000genomes.ebi.ac.uk/vol1/ftp/ on May 31, 2018, IAM Affymetrix 6.0 data was downloaded from ftp.1000genomes.ebi.ac.uk/vol1/ftp/technical/working/20130711_native_american_admix_train on Oct. 11, 2018, ClinVar data was downloaded from https://ftp.ncbi.nlm.nih.gov/pub/clinvar/vcf_GRCh37/ on Oct. 23, 2020. The Summix R package is available at https://github.com/hendriau/Summix and on Bioconductor. The Summix Python package is available at https://github.com/jordanrhall/summix_py.

## References

1. Karczewski, K.J., Francioli, L.C., Tiao, G., Cummings, B.B., Alföldi, J., Wang, Q., Collins, R.L., Laricchia, K.M., Ganna, A., Birnbaum, D.P., et al. (2020). The mutational constraint spectrum quantified from variation in 141,456 humans. Nature 581, 434–443.

2. Karczewski, K.J., Francioli, L.C., Tiao, G., and Cummings, B.B. (2019). Variation across 141,456 human exomes and genomes reveals the spectrum of loss-of-function intolerance across human protein-coding genes. BioRxiv.

3. Phan, L., Jin, Y., Zhang, H., Qiang, W., Shekhtman, E., Shao, D., Revoe, D., Villamarin, R., Ivanchenko, E., Kimura, M., et al. (2020). ALFA: Allele Frequency Aggregator. National Center for Biotechnology Information, US National Library of Medicine. Available Online: www.Ncbi.Nlm.Nih.Gov/snp/docs/gsr/alfa/ (accessed on 10 March 2020).

4. Guo, M.H., Plummer, L., Chan, Y.-M., Hirschhorn, J.N., and Lippincott, M.F. (2018). Burden Testing of Rare Variants Identified through Exome Sequencing via Publicly Available Control Data. Am. J. Hum. Genet. 103, 522–534.

5. Hendricks, A.E., Billups, S.C., Pike, H.N.C., Farooqi, I.S., Zeggini, E., Santorico, S.A., Barroso, I., and Dupuis, J. (2018). ProxECAT: Proxy External Controls Association Test. A new case-control gene region association test using allele frequencies from public controls. PLoS Genet. 14, e1007591.

6. Lee, S., Kim, S., and Fuchsberger, C. (2017). Improving power for rare-variant tests by integrating external controls. Genet. Epidemiol. 41, 610–619.

7. Malki, L., Sarig, O., Romano, M.-T., Méchin, M.-C., Peled, A., Pavlovsky, M., Warshauer, E., Samuelov, L., Uwakwe, L., Briskin, V., et al. (2019). Variant PADI3 in Central Centrifugal Cicatricial Alopecia. N. Engl. J. Med. 380, 833–841.

8. Taliun, D., Harris, D.N., Kessler, M.D., and Carlson, J. (2019). Sequencing of 53,831 diverse genomes from the NHLBI TOPMed Program. BioRxiv.

9. Rannala, B., and Mountain, J.L. (1997). Detecting immigration by using multilocus genotypes. Proc. Natl. Acad. Sci. U. S. A. 94, 9197–9201.

10. Pritchard, J.K., Stephens, M., and Donnelly, P. (2000). Inference of population structure using multilocus genotype data. Genetics 155, 945–959.

11. Tang, H., Peng, J., Wang, P., and Risch, N.J. (2005). Estimation of individual admixture: analytical and study design considerations. Genet. Epidemiol. 28, 289–301.

12. Alexander, D.H., Novembre, J., and Lange, K. (2009). Fast model-based estimation of ancestry in unrelated individuals. Genome Res. 19, 1655–1664.

13. Raj, A., Stephens, M., and Pritchard, J.K. (2014). fastSTRUCTURE: variational inference of population structure in large SNP data sets. Genetics 197, 573–589.

14. Chiang, C.W.K., Gajdos, Z.K.Z., Korn, J.M., Kuruvilla, F.G., Butler, J.L., Hackett, R., Guiducci, C., Nguyen, T.T., Wilks, R., Forrester, T., et al. (2010). Rapid assessment of genetic ancestry in populations of unknown origin by genome-wide genotyping of pooled samples. PLoS Genet. 6, e1000866.

15. Bansal, V., and Libiger, O. (2015). Fast individual ancestry inference from DNA sequence data leveraging allele frequencies for multiple populations. BMC Bioinformatics 16, 4.

16. Shringarpure, S.S., Bustamante, C.D., Lange, K., and Alexander, D.H. (2016). Efficient analysis of large datasets and sex bias with ADMIXTURE. BMC Bioinformatics 17, 218.

17. Sirugo, G., Williams, S.M., and Tishkoff, S.A. (2019). The Missing Diversity in Human Genetic Studies. Cell 177, 26–31.

18. Martin, A.R., Gignoux, C.R., Walters, R.K., Wojcik, G.L., Neale, B.M., Gravel, S., Daly, M.J., Bustamante, C.D., and Kenny, E.E. (2017). Human Demographic History Impacts Genetic Risk Prediction across Diverse Populations. Am. J. Hum. Genet. 100, 635–649.

19. Bonnans, J.-F., Gilbert, J.C., Lemarechal, C., and Sagastizábal, C.A. (2006). Numerical Optimization: Theoretical and Practical Aspects (Springer Science & Business Media).

20. 1000 Genomes Project Consortium, Auton, A., Brooks, L.D., Durbin, R.M., Garrison, E.P., Kang, H.M., Korbel, J.O., Marchini, J.L., McCarthy, S., McVean, G.A., et al. (2015). A global reference for human genetic variation. Nature 526, 68–74.

21. Mao, X., Bigham, A.W., Mei, R., Gutierrez, G., Weiss, K.M., Brutsaert, T.D., Leon-Velarde, F., Moore, L.G., Vargas, E., McKeigue, P.M., et al. (2007). A genomewide admixture mapping panel for Hispanic/Latino populations. Am. J. Hum. Genet. 80, 1171–1178.

22. Wojcik, G.L., Fuchsberger, C., Taliun, D., Welch, R., Martin, A.R., Shringarpure, S., Carlson, C.S., Abecasis, G., Kang, H.M., Boehnke, M., et al. (2018). Imputation-Aware Tag SNP Selection To Improve Power for Large-Scale, Multi-ethnic Association Studies. G3 8, 3255–3267.

23. Kalia, S.S., Adelman, K., Bale, S.J., Chung, W.K., Eng, C., Evans, J.P., Herman, G.E., Hufnagel, S.B., Klein, T.E., Korf, B.R., et al. (2017). Recommendations for reporting of secondary findings in clinical exome and genome sequencing, 2016 update (ACMG SF v2.0): a policy statement of the American College of Medical Genetics and Genomics. Genet. Med. 19, 249–255.

24. Bhérer, C., Campbell, C.L., and Auton, A. (2017). Refined genetic maps reveal sexual dimorphism in human meiotic recombination at multiple scales. Nat. Commun. 8, 14994.

25. Zeileis, A., and Grothendieck, G. zoo: S3 Infrastructure for Regular and Irregular Time Series.

26. Bates, D., Mächler, M., Bolker, B., and Walker, S. (2015). Fitting Linear Mixed-Effects Models Usinglme4. Journal of Statistical Software 67,.

27. Watson, P.F., and Petrie, A. (2010). Method agreement analysis: a review of correct methodology. Theriogenology 73, 1167–1179.

28. Lin, L.I. (1989). A concordance correlation coefficient to evaluate reproducibility. Biometrics 45, 255–268.

29. Signorell, A., Aho, K., Anderegg, N., Aragon, T., Arppe, A., Baddeley, A., Bolker, B., Caeiro, F., Champely, S., Chessel, D., et al. (2018). DescTools: Tools for descriptive statistics. 2018. R Package Version 0. 99 24,.

30. Nakatsuka, N., Moorjani, P., Rai, N., Sarkar, B., Tandon, A., Patterson, N., Bhavani, G.S., Girisha, K.M., Mustak, M.S., Srinivasan, S., et al. (2017). The promise of discovering population-specific disease - associated genes in South Asia. Nat. Genet. 49, 1403–1407.

31. Narasimhan, V.M., Patterson, N., Moorjani, P., Rohland, N., Bernardos, R., Mallick, S., Lazaridis, I., Nakatsuka, N., Olalde, I., Lipson, M., et al. (2019). The formation of human populations in South and Central Asia. Science 365,.

32. Reich, D., Thangaraj, K., Patterson, N., Price, A.L., and Singh, L. (2009). Reconstructing Indian population history. Nature 461, 489–494.

33. Richards, S., Aziz, N., Bale, S., Bick, D., Das, S., Gastier-Foster, J., Grody, W.W., Hegde, M., Lyon, E., Spector, E., et al. (2015). Standards and guidelines for the interpretation of sequence variants: a joint consensus recommendation of the American College of Medical Genetics and Genomics and the Association for Molecular Pathology. Genet. Med. 17, 405–423.

34. Farrer, L.A., Cupples, L.A., Haines, J.L., Hyman, B., Kukull, W.A., Mayeux, R., Myers, R.H., Pericak-Vance, M.A., Risch, N., and Van Duijn, C.M. (1997). APOE and Alzheimer Disease Meta Analysis Consortium Effects of age, sex, and ethnicity on the association between apolipoprotein E genotype and Alzheimer disease. A meta-analysis. JAMA 278, 1349–1356.

35. Graff-Radford, N.R., Green, R.C., Go, R.C.P., Hutton, M.L., Edeki, T., Bachman, D., Adamson, J.L., Griffith, P., Willis, F.B., Williams, M., et al. (2002). Association between apolipoprotein E genotype and Alzheimer disease in African American subjects. Arch. Neurol. 59, 594–600.

36. Logue, M.W., Schu, M., Vardarajan, B.N., Buros, J., Green, R.C., Go, R.C.P., Griffith, P., Obisesan, T.O., Shatz, R., Borenstein, A., et al. (2011). A comprehensive genetic association study of Alzheimer disease in African Americans. Arch. Neurol. 68, 1569–1579.

37. Blue, E.E., Horimoto, A.R.V.R., Mukherjee, S., Wijsman, E.M., and Thornton, T.A. (2019). Local ancestry at APOE modifies Alzheimer’s disease risk in Caribbean Hispanics. Alzheimers. Dement. 15, 1524–1532.

38. Montana, G., and Pritchard, J.K. (2004). Statistical tests for admixture mapping with case-control and cases-only data. Am. J. Hum. Genet. 75, 771–789.

39. Zhou, Q., Zhao, L., and Guan, Y. (2016). Strong Selection at MHC in Mexicans since Admixture. PLoS Genet. 12, e1005847.

40. Hodgson, J.A., Pickrell, J.K., Pearson, L.N., Quillen, E.E., Prista, A., Rocha, J., Soodyall, H., Shriver, M.D., and Perry, G.H. (2014). Natural selection for the Duffy-null allele in the recently admixed people of Madagascar. Proc. Biol. Sci. 281, 20140930.

41. Brown, R., and Pasaniuc, B. (2014). Enhanced methods for local ancestry assignment in sequenced admixed individuals. PLoS Comput. Biol. 10, e1003555.

42. Jiang, L., Jiang, H., Dai, S., Chen, Y., Song, Y., Tang, C.S.-M., Wang, B., Garcia-Barcelo, M.-M., Tam, P., Cherny, S.S., et al. (2020). Deviation from baseline mutation burden provides powerful and robust rare - variants association test for complex diseases.

43. Li, Y., and Lee, S. (2020). Novel score test to increase power in association test by integrating external controls. Genet. Epidemiol.

44. Buniello, A., MacArthur, J.A.L., Cerezo, M., Harris, L.W., Hayhurst, J., Malangone, C., McMahon, A., Morales, J., Mountjoy, E., Sollis, E., et al. (2019). The NHGRI-EBI GWAS Catalog of published genome - wide association studies, targeted arrays and summary statistics 2019. Nucleic Acids Res. 47, D1005–D1012.

45. Gong, T., and Szustakowski, J.D. (2013). DeconRNASeq: a statistical framework for deconvolution of heterogeneous tissue samples based on mRNA-Seq data. Bioinformatics 29, 1083–1085.

46. Hao, Y., Yan, M., Heath, B.R., Lei, Y.L., and Xie, Y. (2019). Fast and robust deconvolution of tumor infiltrating lymphocyte from expression profiles using least trimmed squares. PLoS Comput. Biol. 15, e1006976.

47. Racle, J., de Jonge, K., Baumgaertner, P., Speiser, D.E., and Gfeller, D. (2017). Simultaneous enumeration of cancer and immune cell types from bulk tumor gene expression data. Elife 6,.

