## Supplementary material for "*Summix:* A method for detecting and adjusting for population structure in genetic summary data": Figures S1-S11, Tables S12-S13

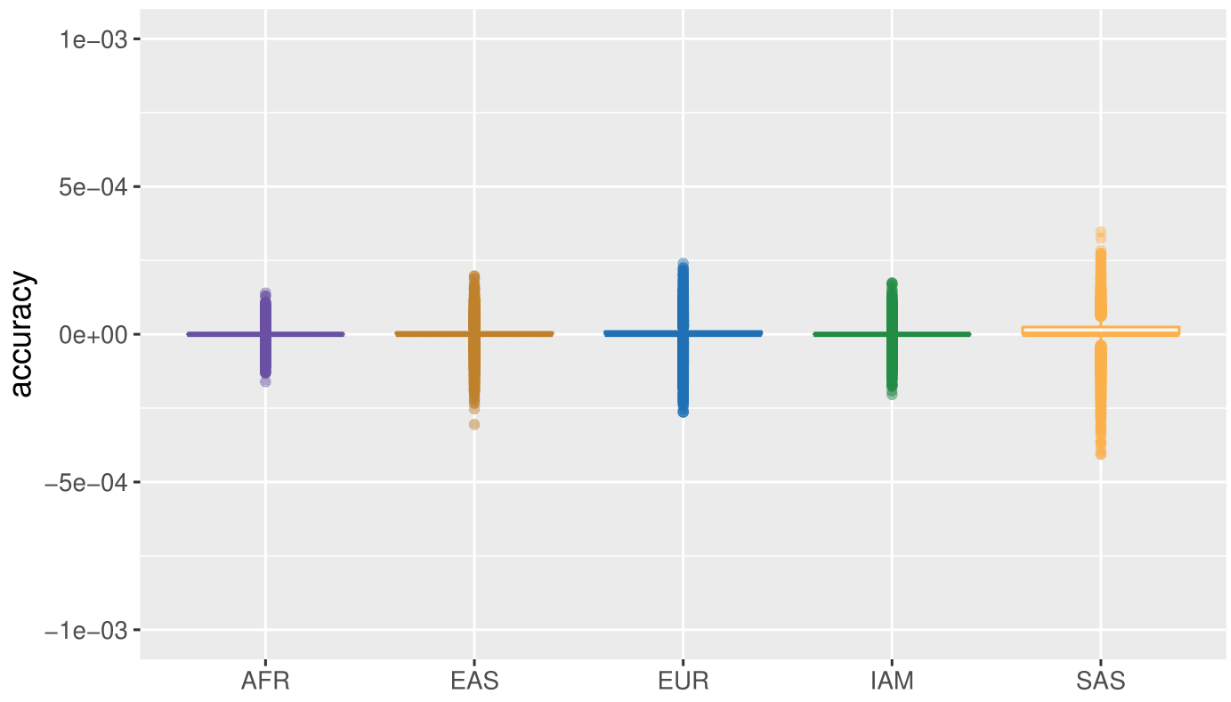

**Figure S1. Simulation results for one ancestry parameters**. Accuracy is defined as the absolute difference between the estimated ancestry proportions and given ancestry proportions within simulations. A single reference ancestry was used to simulate genotypes of a population.

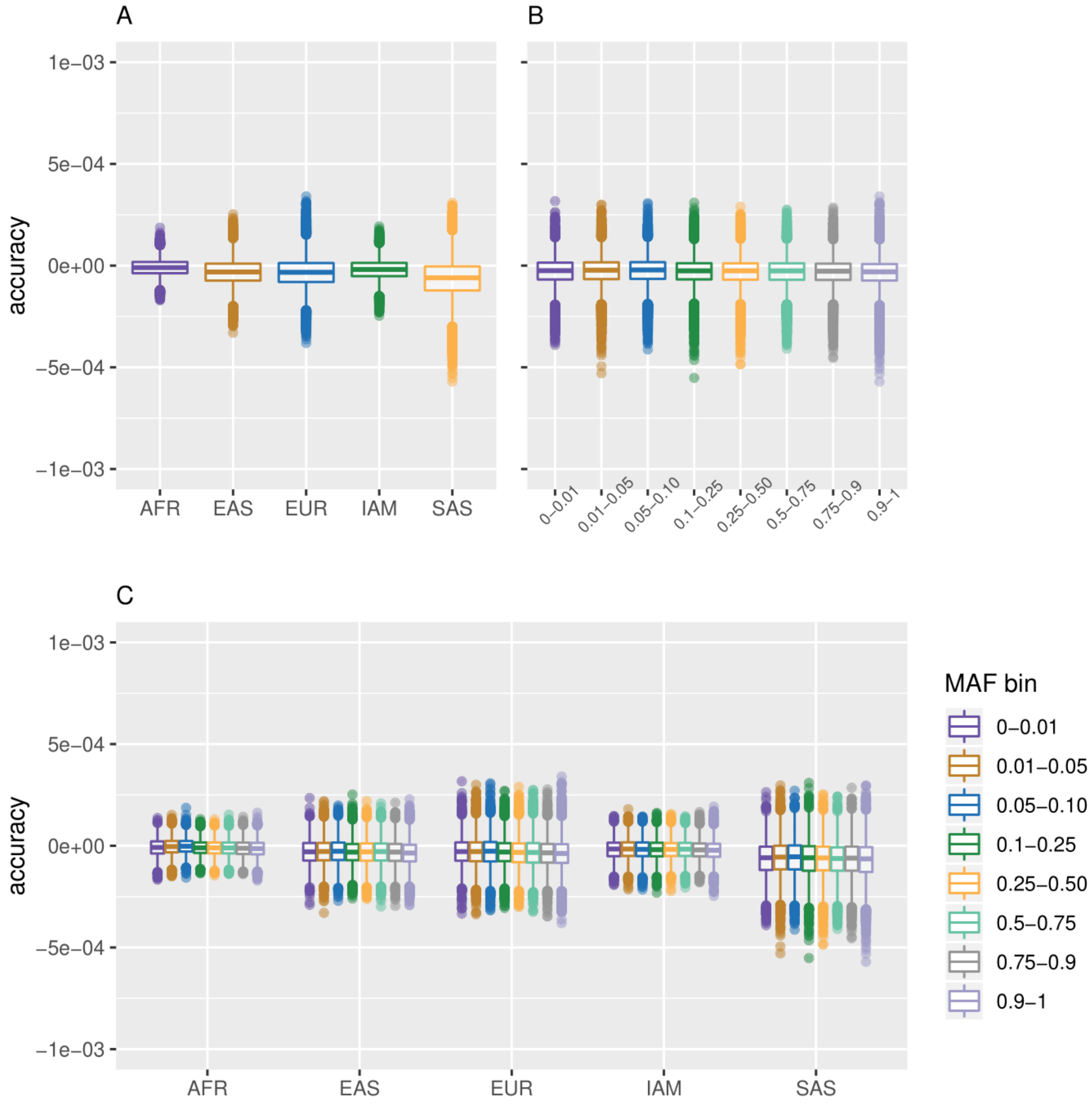

**Figure S2. Simulation results for two ancestry parameters**. Accuracy is defined as the absolute difference between the estimated ancestry proportions and given ancestry proportions within simulations. Two reference ancestries were used to simulate genotypes of an admixed population. **A)** Accuracy separated by ancestry. **B)** Accuracy separated by minor allele frequency. **C)** Accuracy separated by both ancestry and minor allele frequency.

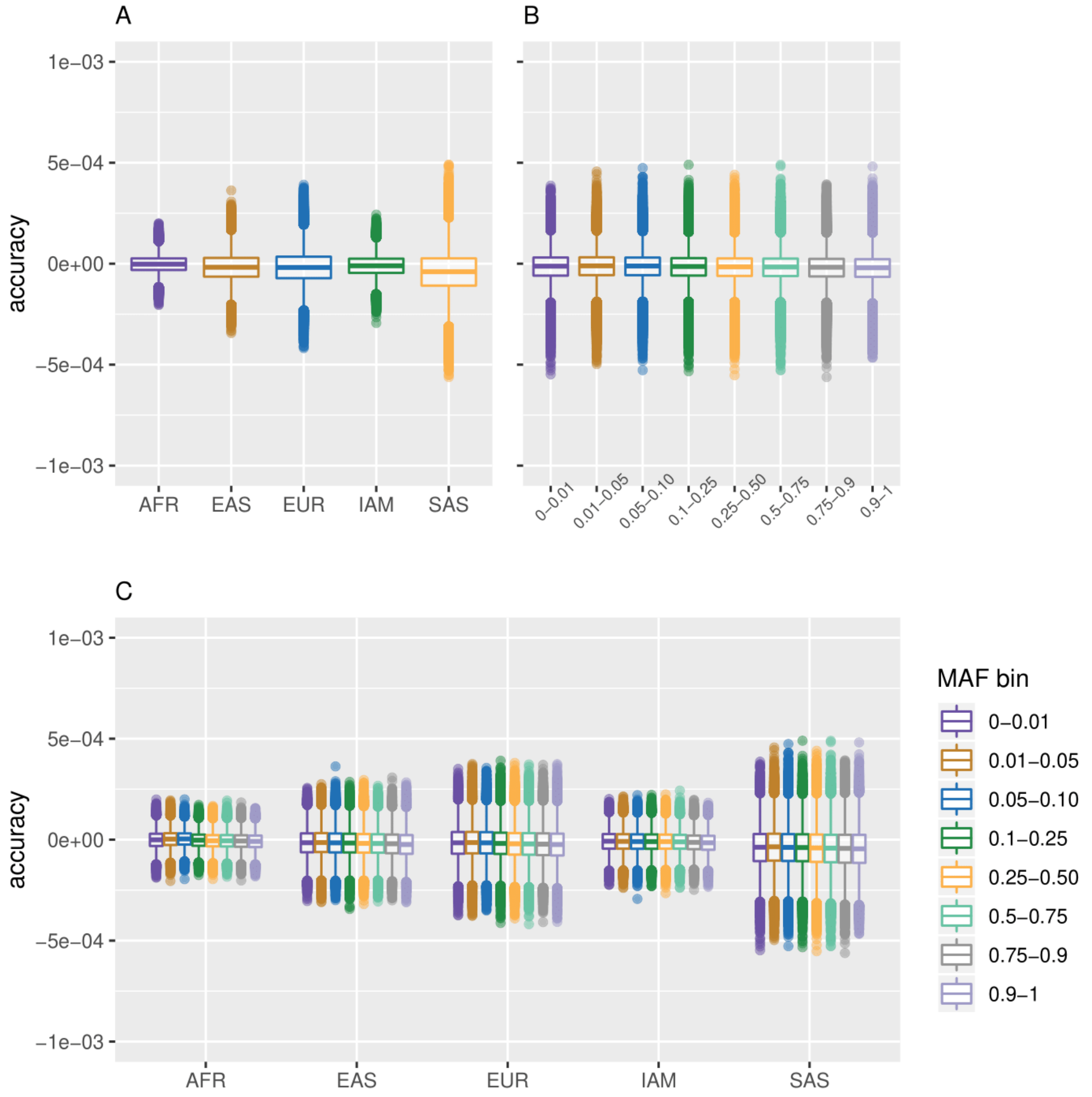

**Figure S3. Simulation results for three ancestry parameters**. Accuracy is defined as the absolute difference between the estimated ancestry proportions and given ancestry proportions within simulations. Three reference ancestries were used to simulate genotypes of an admixed population. **A)** Accuracy separated by ancestry. **B)** Accuracy separated by minor allele frequency. **C)** Accuracy separated by both ancestry and minor allele frequency.

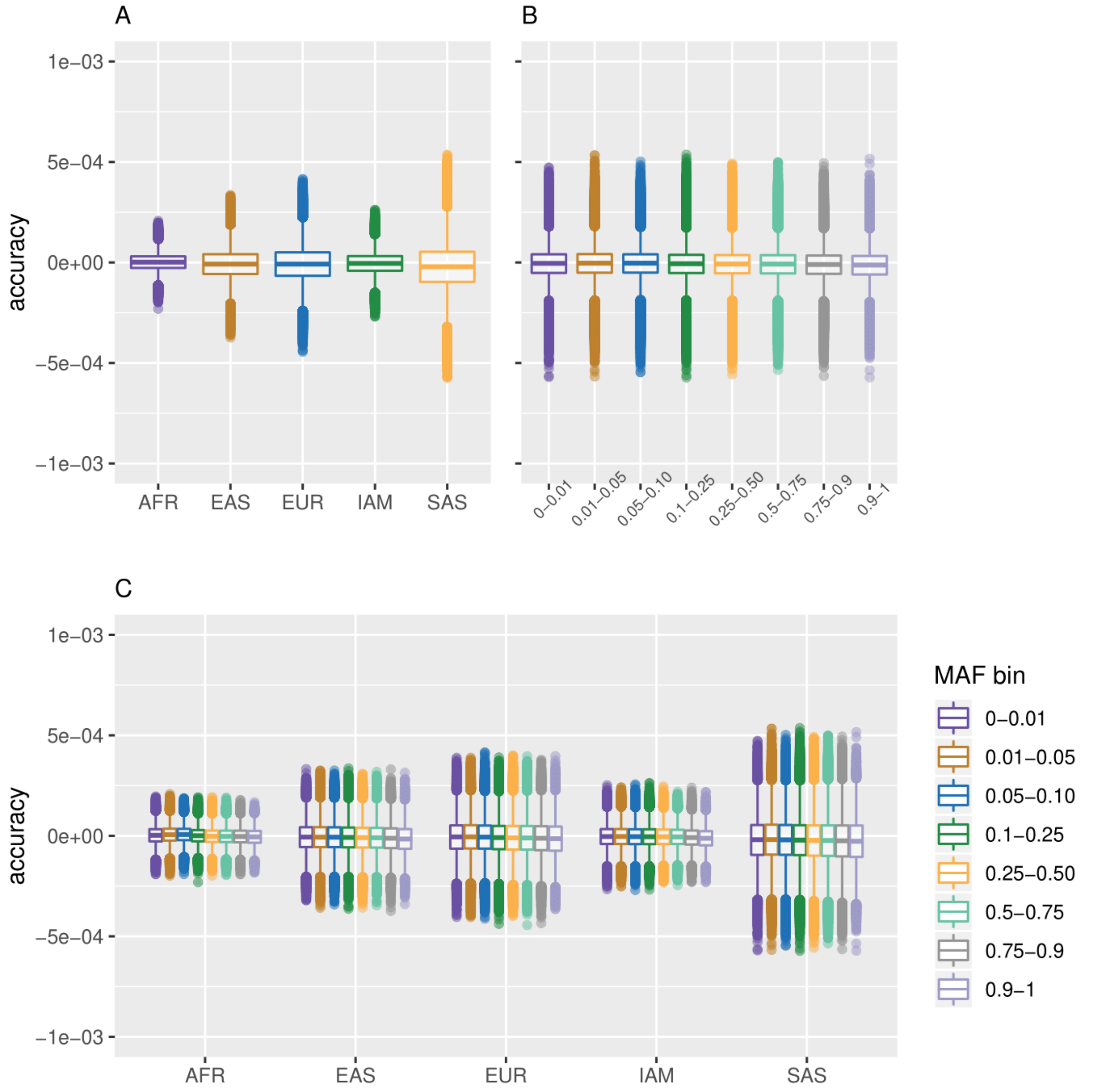

**Figure S4. Simulation results for four ancestry parameters**. Accuracy is defined as the absolute difference between the estimated ancestry proportions and given ancestry proportions within simulations. Four reference ancestries were used to simulate genotypes of an admixed population. **A)** Accuracy separated by ancestry. **B)** Accuracy separated by minor allele frequency. **C)** Accuracy separated by both ancestry and minor allele frequency.

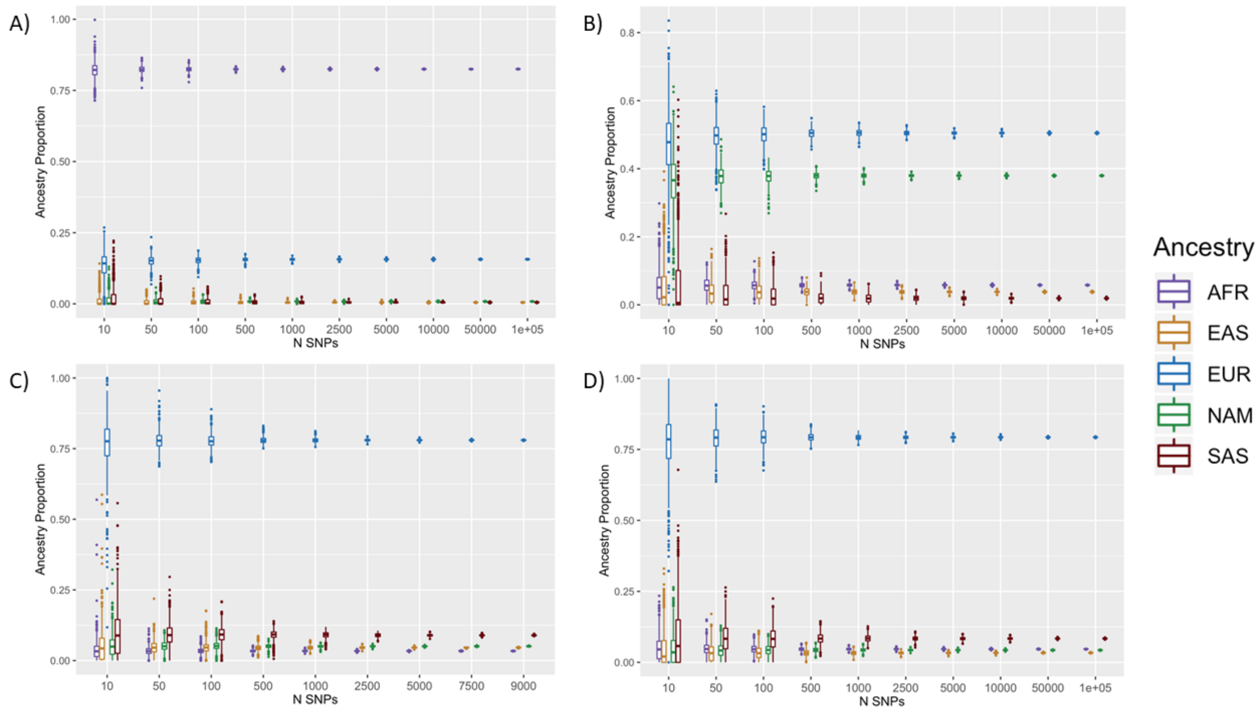

**Figure S5. Precision in ancestry estimates for AFR, AMR and OTH gnomad groups by number of SNPs.** Number of SNPs (x-axis), estimated ancestry proportion (y-axis) for 1,000 replicates; **A)** AFR genome. **B)** AMR genome. **C)** OTH exome. **D)** OTH genome.

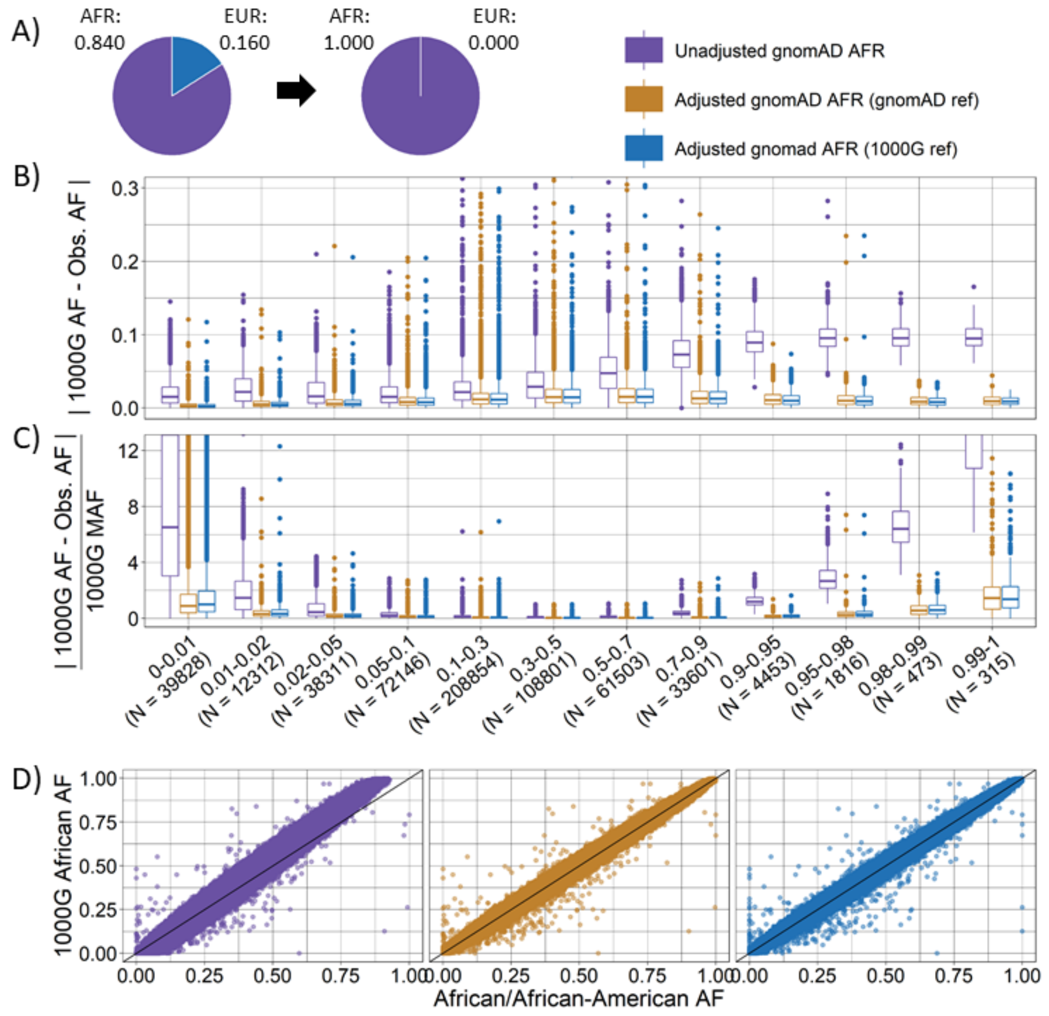

**Figure S6. Ancestry-adjusted vs. unadjusted allele frequency for gnomAD African/African American genomes for a target sample with African ancestry.** Ancestry-adjusted AF was estimated for a target sample with 100% African ancestry using gnomAD (orange) or 1000 Genomes (blue) Non-Finnish European as reference and compared to unadjusted AF (purple) for 582,413 SNPs. **A)** ancestry proportions for gnomAD African/African American gnomes (AFR = 0.840, EUR=0.160) and target sample (AFR = 1); **B)** absolute difference between target sample AF (1000 Genomes African ancestry) and unadjusted or ancestry-adjusted gnomAD AF by 1000 Genomes AF category; **C)** relative difference between target 1000 Genomes African ancestry AF and unadjusted or ancestry-adjusted gnomAD AF by 1000 Genomes AF category; unzoomed figures B and C are available in the supplemental (**Figure S7**). **D)** scatter plot of target sample 1000 Genomes AF (y-axis) and unadjusted (left), ancestry-adjusted with gnomAD reference (center), and ancestry-adjusted with 1000 Genomes reference (right) gnomAD AF (x-axis).

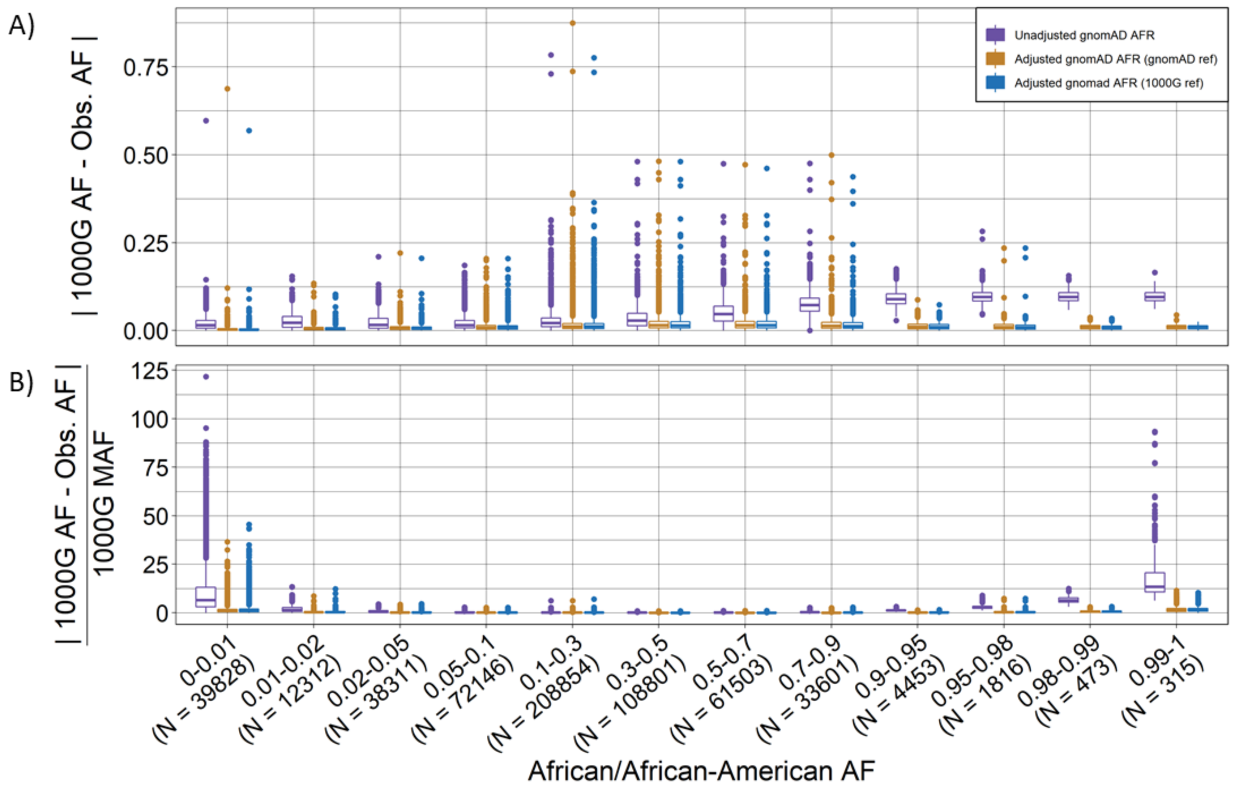

**Figure S7. Unzoomed Ancestry-adjusted vs. unadjusted allele frequency for gnomAD African/African American genomes for a target sample with African ancestry (Complimentary to Figure S6). A)** absolute difference between target sample AF (1000 Genomes African ancestry) and unadjusted or ancestry-adjusted gnomAD AF by 1000 Genomes AF category; **B)** relative difference between target 1000 Genomes African ancestry AF and unadjusted or ancestry-adjusted gnomAD AF by 1000 Genomes AF category.

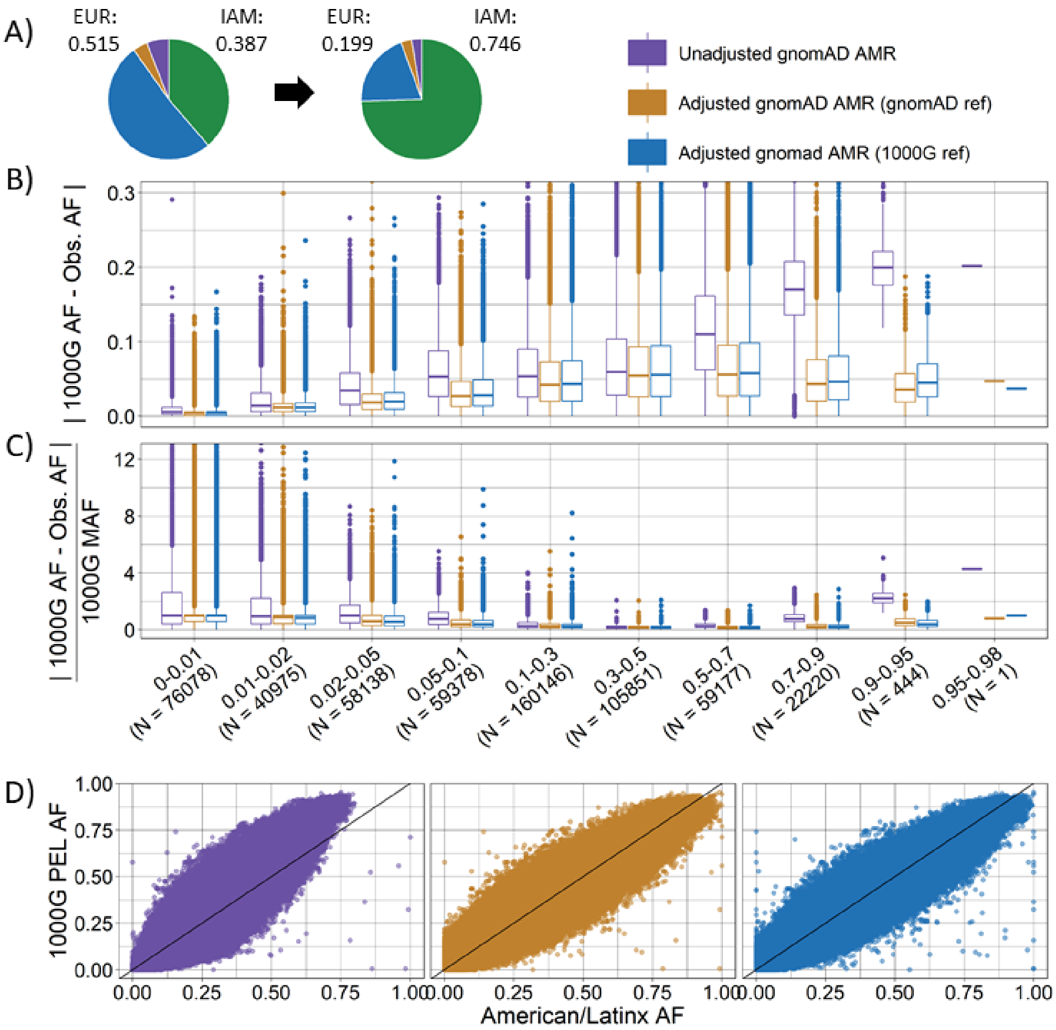

**Figure S8. Ancestry-adjusted vs. unadjusted allele frequency for gnomAD American/Latinx genomes for a target sample of Peruvian ancestry.** Ancestry-adjusted AF was estimated for a target Peruvian sample using gnomAD (orange) or 1000 Genomes (blue) East Asian, European, and African as reference ancestral populations and compared to unadjusted AF (purple) for 582,408 SNPs. **A)** normalized ancestry proportions estimated for gnomAD American/Latinx genomes (purple AFR = 0.059, orange EAS=0.039, blue EUR=0.387, green IAM=0.515) and target Peruvian ancestry proportions (purple AFR = 0.028, orange EAS=0.027, blue EUR=0.199, green IAM=0.746); **B)** absolute difference between target 1000 Genomes Peruvian ancestry AF and unadjusted or ancestry-adjusted gnomAD AF by 1000 Genomes AF category; **C)** relative difference between target 1000 Genomes Peruvian ancestry AF and unadjusted or ancestry-adjusted gnomAD AF by 1000 Genomes AF category; unzoomed figures B and C are available in the supplemental (**Figure S9**). **D)** scatter plot of target 1000 Genomes AF (y-axis) and unadjusted (left), ancestry-adjusted with gnomAD reference (center), and ancestry-adjusted with 1000 Genomes reference (right) gnomAD AF (x-axis).

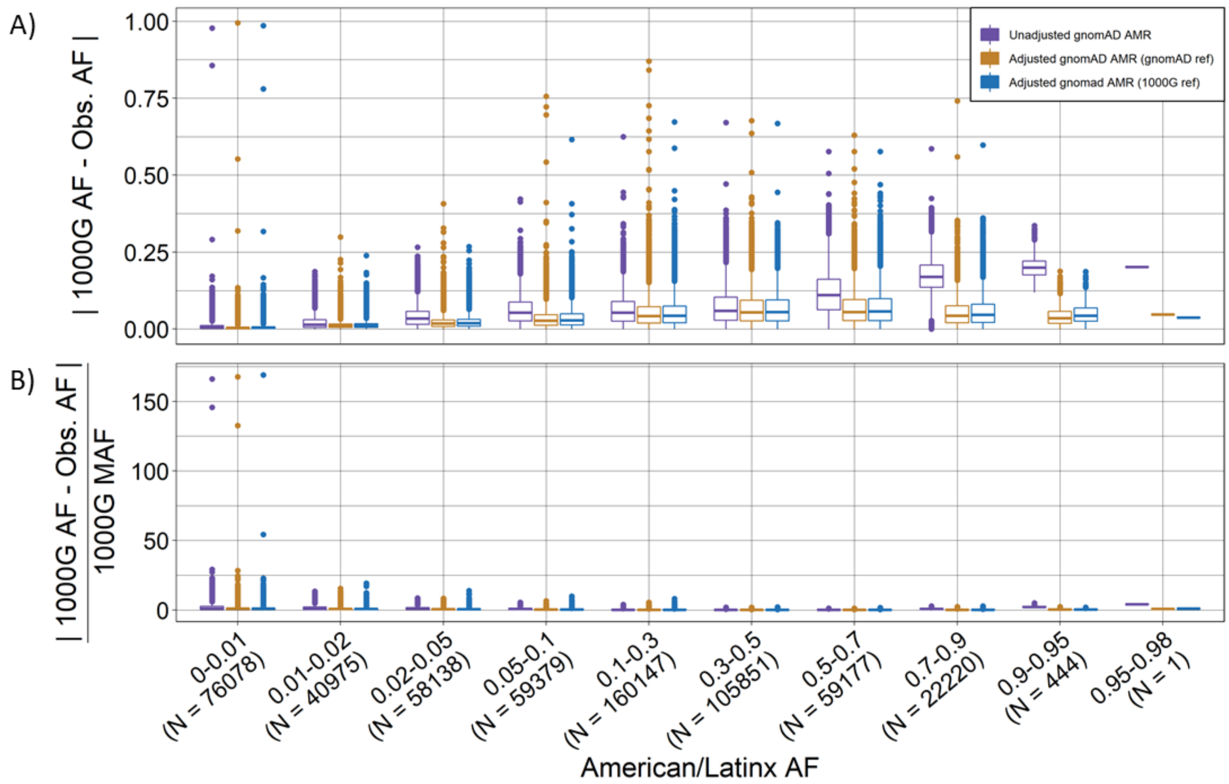

**Figure S9. Unzoomed Ancestry-adjusted vs. unadjusted allele frequency for gnomAD American/Latinx genomes for a target sample of Peruvian ancestry (Complimentary to Figure S8). A)** absolute difference between target 1000 Genomes Peruvian ancestry AF and unadjusted or ancestry-adjusted gnomAD AF by 1000 Genomes AF category; **B)** relative difference between target 1000 Genomes Peruvian ancestry AF and unadjusted or ancestry-adjusted gnomAD AF by 1000 Genomes AF category.

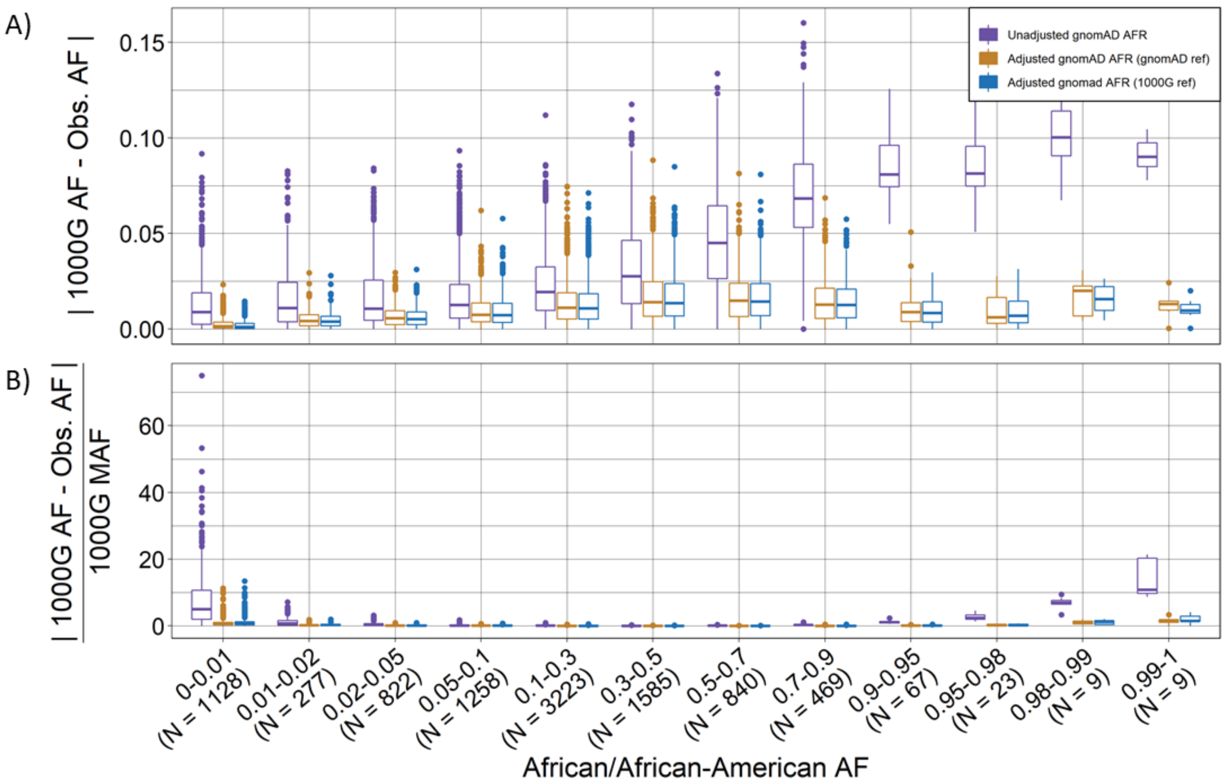

**Figure S10. Unzoomed Ancestry-adjusted vs. unadjusted allele frequency for gnomAD African/African American genomes for a target sample with African ancestry (Complimentary to Figure 3). A)** absolute difference between target sample AF (1000 Genomes African ancestry) and unadjusted or ancestry-adjusted gnomAD AF by 1000 Genomes AF category; **B)** relative difference between target 1000 Genomes African ancestry AF and unadjusted or ancestry-adjusted gnomAD AF by 1000 Genomes AF category.

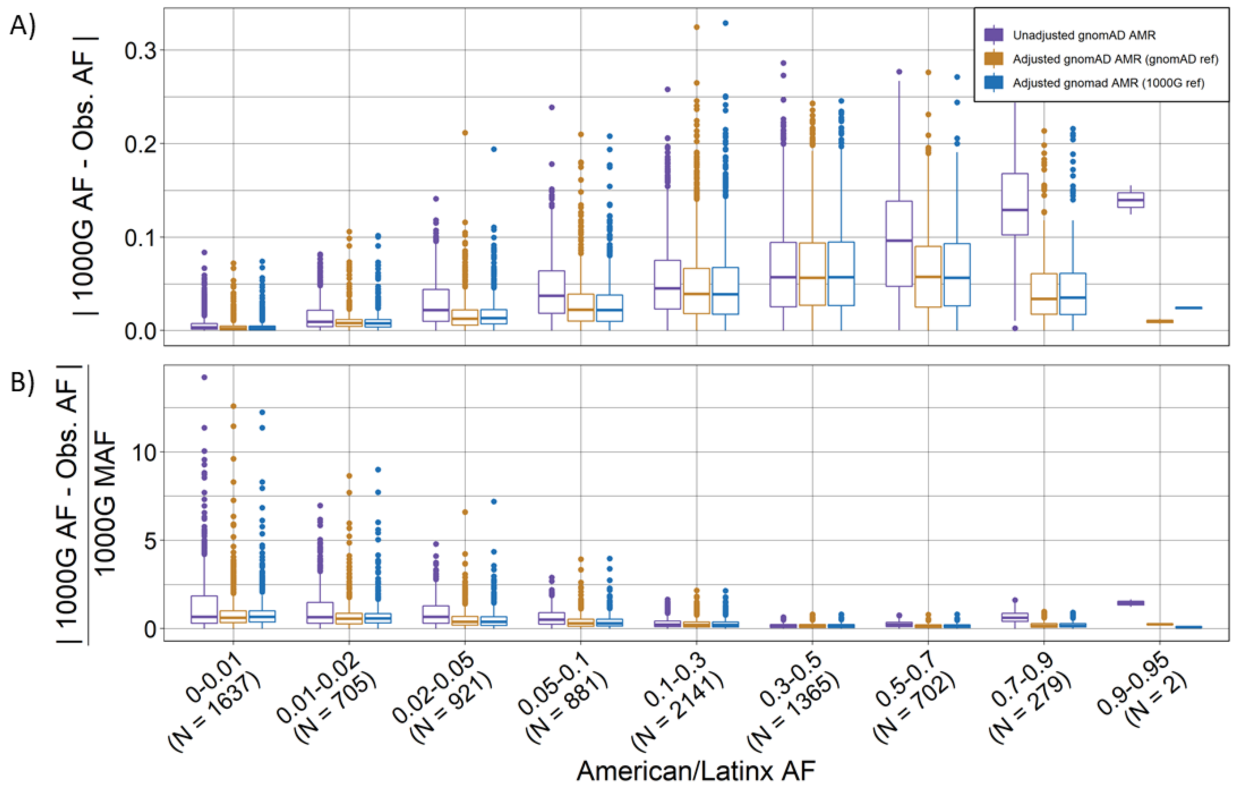

**Figure S11. Unzoomed Ancestry-adjusted vs. unadjusted allele frequency for gnomAD American/Latinx exomes for a target sample of Peruvian ancestry (Complimentary to Figure 4). A)** absolute difference between target 1000 Genomes Peruvian ancestry AF and unadjusted or ancestry-adjusted gnomAD AF by 1000 Genomes AF category; **B)** relative difference between target 1000 Genomes Peruvian ancestry AF and unadjusted or ancestry-adjusted gnomAD AF by 1000 Genomes AF category.

**Table S12**. Unadjusted and adjusted values for *PADI3* variants from Malki et al. 2019.

| rsID<br>Location<br>DNA Sequence Variant<br>AA Sequence Change | AN | unadjusted | | | 100% AFR adjusted | | |
|---|---|---|---|---|---|---|---|
| | | AF | AC | Number of Homozygotes | AF | AC* | Number of Homozygotes** |
| rs139426141<br>1-17597398-A-G<br>c.856A→G<br>p.Thr286Ala | 24952 | 0.0365 | 910 | 23 | 0.0434 | 1082 | 47 |
| rs34097903<br>1-17607274-G-A<br>c.1744G→A<br>p.Ala582Thr | 24964 | 0.0227 | 566 | 6 | 0.0270 | 673 | 18 |
| rs140482516<br>1-17607199-C-T<br>c.1669C→T<br>p.Arg557Trp | 24968 | 0.0075 | 188 | 0 | 0.0089 | 223 | 2 |
| rs1557508308<br>1-17597372-A-G<br>c.832-2A→G<br>splicing | 0 | 0 | 0 | 0 | 0 | 0 | 0 |
| rs1437225536<br>1-17609534-G-A<br>c.1955G→A<br>p.Arg652Lys | 8716 | 0.0001 | 1 | 0 | 0.0001 | 1 | 0 |
| rs139876092<br>1-17594433-C-T<br>c.628C→T<br>p.Arg210Trp | 24962 | 0.00164 | 41 | 0 | 0.0019 | 48 | 0 |
| Total | -- | -- | 1706 | 29 | -- | 2027 | 67 |

* $AC_{adj} = round(AF_{adj} * AN)$

**Adjusted number of homozygotes was estimated assuming Hardy-Weinberg equilibrium, $N_{homozygotes} = round(AF_{adj}^2 * AN)$

**Table S13**. Number of cases and African/African American gnomAD v2.1 controls with minor alleles in *PADI3* variants reported from Malki et al.

| | Cases | gnomAD v2.1 African | |
|---|---|---|---|
| | | unadjusted | 100% AFR adjusted*** |
| Minor allele | 14 | 1677* | 1960 |
| No minor allele | 44 | 10810** | 10527 |
| Chi Square p-value | | 0.029 | 0.114 |
| Fisher's exact test p-value | | 0.031 | 0.101 |

* number of individuals with at least one minor allele was estimated by removing the number of homozygotes from the total minor allele count

** number of individuals with no minor allele was estimated as the total number of gnomAD v2.1 African/African American individuals (N=12,487) minus the number of estimates individuals with at least one minor allele

*** adjusted number of individuals was estimated by $\sum_{j=1}^{K} round\left(AN_j * AF_{adj_j} - AN_j * AF_{adj_j}^2\right)$ where $AN_j$ and $AF_{adj_j}$ are the observed allele number and adjusted allele frequency for variant j respectively. The values are rounded to the nearest integer before summing.
